# Lamin B1 Safeguards the B Cell Genome and Shapes Lymphoma Outcome

**DOI:** 10.1101/2025.11.03.686246

**Authors:** Filip Filipsky, Katarina B Chapman, Johannes Bloehdorn, Jun Wang, John Gribben, Michael Hausmann, Christoph Cremer, Andrejs Braun, Marta C Sallan, Tanya Klymenko

## Abstract

Lamin B1 is a structural component of the nuclear lamina that participates in diverse cellular processes, including genome regulation and cellular senescence. During adaptive immune responses, B lymphocytes in germinal centres (GCs) undergo clonal expansion and programmed DNA damage at immunoglobulin loci, while simultaneously downregulating Lamin B1. Likewise, Lamin B1 downregulation has been observed in GC-derived lymphomas and myeloid malignancies, yet the functional consequences of Lamin B1 loss during B cell development remain poorly understood. Here, we used *in vivo* and *in vitro* B cell models of conditional hypomorphic expression of Lamin B1, which showed elevated DNA damage, altered chromatin accessibility, and disrupted transcriptional profiles. Using sBLISS (*in situ* labelling and sequencing of double-strand breaks), we identified non-random double-strand break hotspots in both mouse and human GC B cells, depleted of Lamin B1. These breaks are preferentially located near transcriptional start sites (TSSs) and regulatory elements that control translation and mRNA fate, implicating Lamin B1 in protecting fragile regulatory regions. Moreover, low LMNB1 expression correlated with poor clinical outcomes in patients with diffuse large B cell lymphoma (DLBCL). Together, this study reveals a crucial role for Lamin B1 in preserving genomic stability in B cells, underscoring its impact on the pathogenesis of B cell-derived malignancies.

## Introduction

The nuclear lamina (NL) is a cellular component that contributes to a wide range of biological processes, including genome organisation, gene transcription, and mechanical support by stabilising the nuclear envelope (Misteli, 2020). The NL comprises type V intermediate filaments (Lamin A/C, Lamin B1/ B2), which are in close contact with the nuclear membrane and chromatin (Aebi, Cohn et al., 1986, Shimi, Pfleghaar et al., 2008). Together with chromatin, Lamin B1 forms genomic regions known as lamina-associated domains (LADs) (Guelen, Pagie et al., 2008). The removal of Lamin B1 from the nuclear periphery prompts major conformational changes in LADs during differentiation, leading to chromatin reorganisation and differential gene expression (Sadaie, Salama et al., 2013, Shah, Donahue et al., 2013). Due to the essential role of Lamin B1 in various cellular functions, there is limited evidence of pathogenesis linked directly to the dysregulation of Lamin B1, except in autosomal dominant leukodystrophy (ADLD) (Padiath, Saigoh et al., 2006). and acquired Pelger–Huët Anomaly (PHA) (Reilly, Philip Creamer et al., 2022).

Constant expression of Lamin B1 within the nucleus likely reflects low protein turnover and high stability caused by extensive post-translational modifications (Adam, Butin-Israeli et al., 2013, Jung, Nobumori et al., 2013, Simon & Wilson, 2013). In the last decade, a number of empirical studies have demonstrated that Lamin B1 might play a role in malignant transformation and be involved in the clinical outcomes of cancer.

However, the potential clinical predictive value of Lamin B1 is context-dependent and *LMNB1* reduction can be linked to either favourable (Izdebska, Gagat et al., 2018, Li, Du et al., 2013, Qin, Lu et al., 2022) or poor clinical outcomes (Klymenko, Bloehdorn et al., 2018, Moss, Krivosheyev et al., 1999, Reilly et al., 2022). Within the physiological context of the B cell development, Lamin B1 loss in haematopoietic stem cells (HSC) is responsible for abnormal nuclear morphology followed by dysfunctional progenitor cell fate determination via genome organisation (Reilly et al., 2022). In cell-mediated immune responses, Lamin B1 is functionally downregulated at the nuclear periphery in germinal centre (GC) B cells, leading to increased accessibility of the immunoglobulin variable region (IgV) (Klymenko et al., 2018), suggesting a direct role of the nuclear lamina in safeguarding the genome from point mutagenesis.

Although the exact mechanism of lamina-mediated mutagenesis remains unknown, current evidence suggests that disruption of Lamin B1 nuclear localisation might initiate chromatin relaxation and induce the formation of transcriptionally active chromatin (Reddy, Zullo et al., 2008, Sadaie et al., 2013, Shah et al., 2013). While other nuclear lamins (Lamin B2, Lamin A/C) are also in contact with heterochromatin at the nuclear periphery, Lamin B1 is associated with active euchromatin in the nucleoplasm during the epithelial-to-mesenchymal transition (EMT) in mice (Gesson, Rescheneder et al., 2016, Pascual-Reguant, Blanco et al., 2018). In addition to transcriptionally silent LADs, Lamin B1 forms highly dynamic euchromatin LADs (eLADs), further implying complex and precise localisation of Lamin B1 (Pascual-Reguant et al., 2018).

Amongst the documented, multifaceted roles of Lamin B1, previous studies described its involvement in mediating the DNA damage response (DDR) and chromosomal instability (Butin-Israeli, Adam et al., 2013, Butin-Israeli, Adam et al., 2015, Liu, Sun et al., 2015, Murray-Nerger, Justice et al., 2021). Consequently, chromosomal translocations arising from activation-induced cytidine deaminase (AID) mediated DSBs during B cell activation are key genetic events that contribute to lymphomagenesis (Liu, Duke et al., 2008, Nussenzweig & Nussenzweig, 2010). Under physiological conditions, B cell maturation and antibody affinity are dependent on efficient SHM and class-switch recombination (CSR) (Boboila, Alt et al., 2012), where targeted DNA damage during CSR is resolved by the non-homologous end joining (NHEJ) repair pathway (Bothmer, Robbiani et al., 2010, Gupta, Hunt et al., 2014). An upstream mediator of NHEJ is p53-binding protein (53BP1), which, in association with other complexes, protects double-stranded ends and prevents the deployment of alternative repair pathways, such as homologous recombination (HR) (Noordermeer, Adam et al., 2018). Etourneaud et al. (2021) demonstrated that overexpression of Lamin B1 prevents the recruitment of 53BP1 to damaged sites (Etourneaud, Moussa et al., 2021), implying that Lamin B1 has a crucial influence on the efficiency of NHEJ. However, while these *in vitro* findings shed light on the involvement of Lamin B1 in the DDR, its precise role in the context of genomic instability in B cells *in vivo* remains unclear. Likewise, there is currently limited data directly linking lamina-modulated DDR with carcinogenesis.

Here, we assessed the effect of Lamin B1 depletion on genomic instability in GC B cells by using hypomorphic Lamin B1 *in vitro* and *in vivo* experimental models. Complementing the functional studies, we also implemented a multi-omics approach that enabled us to understand how the downregulation of Lamin B1 in B cells affects genomic instability and other cellular processes. Our data provide novel insights into the unique role of Lamin B1 in genomic instability in both normal and malignant B cells. Clinical analyses revealed that Lamin B1 downregulation is associated with poor survival outcomes in DLBCL patients.

## Results

### Decreased Lamin B1 is associated with genomic instability and mutagenesis in leukaemia

To establish a functional *in vivo* link between Lamin B1 expression and genomic instability, we evaluated whether the loss of *LMNB1* correlates with genomic instability in patients with CLL. For this, we first assessed the respective *LMNB1* expression according to consensus clustering analysis and the corresponding genomic categorisation (Bloehdorn, Braun et al., 2021). We observed a significant decrease in *LMNB1* expression, accompanied by elevated levels of DNA repair genes (base excision repair (BER), mismatch repair (MMR)), particularly a subset of NHEJ-related genes (*NHEJ1, RAD50, XRCC4, XRCC5*) in the GI (“genomically instable, non-inflammatory”) classified CLL patients compared to (I)-EMT-L (“epithelial-mesenchymal-transition-like with inflammatory features)” subtype (Fig. 1A-C; Fig. S1A).

**Figure 1.**
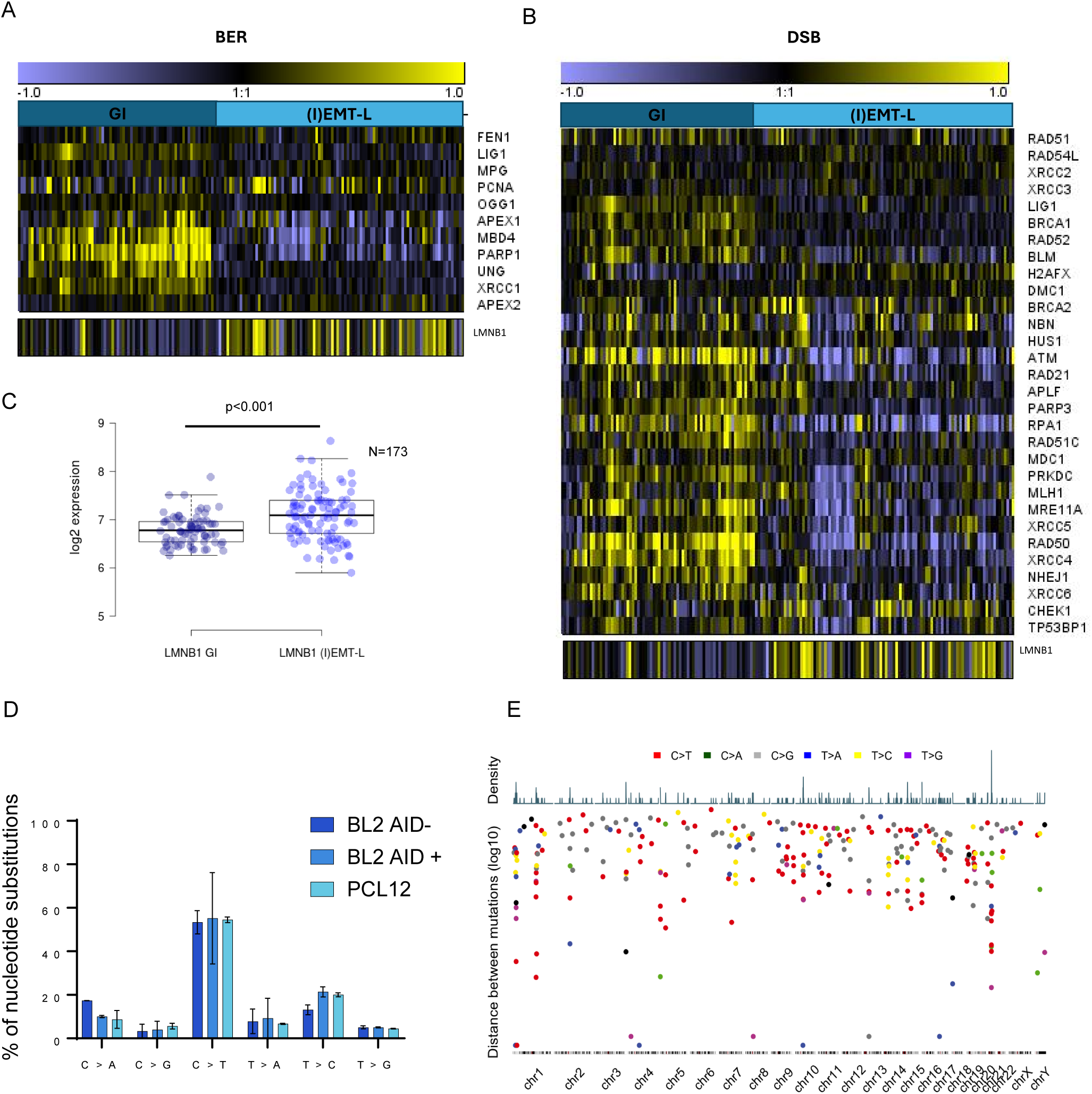
Decreased *LMNB1* expression is associated with genomic instability (GI) in CLL and mutagenesis in malignant B cells. (**A-B**) Heatmaps showing GEP for *LMNB1* and (**A**) base excision repair (BER) and (**B**) DNA double-strand break (DSB) repair genes in GI and (I)EMT-L CLL subtypes (*n* = 173). (**C**) *LMNB1* expression is significantly decreased in GI vs. (I)EMT-L CLL patients. **(D)** Unique somatic variants detected in siLMNB1 samples are represented as the proportion of cytosine to thymine (C>T) nucleotide substitutions. WES was performed on two independent replicates. Error bars represent +/- s.e.m. **(E)** Mutational topology of SNVs detected specifically in siLMNB1-treated BL2 *AID^wt^*, BL2 *AID^-/-^*, and PCL12 cells. Distribution of mutations was plotted in R with the karyoplotR (1.22.0) package, and hg38 was used for the chromosome visualisation.

To identify the Lamin B1-controlled genome regions, we next performed deep whole-exome sequencing (WES) after depleting *LMNB1* in BL2 *AID^wt^*, BL2 *AID*^-/-^, and PCL12 cell lines (Fig. S1B). In Lamin B1-depleted cells, Mutect2 identified single-nucleotide variants (SNVs) consistent with the mutational signature of AID (Alexandrov, Nik-Zainal et al., 2013), where a cytosine-to-thymine (C>T) nucleotide substitution pattern was observed irrespective of AID expression status (Fig. 1D; S1C). Within this context, topological mapping of SNVs revealed genome-wide non-uniform clustering of mutations in Lamin B1-depleted cells (Fig. 1E). A comparison of mutational clusters showed minimal overlap in BL2 *AID^wt^* and *AID^-/-^*samples, further indicating that mutagenesis is coupled with Lamin B1 depletion independent of AID expression. These findings demonstrate that decreased Lamin B1 expression is associated with genomic instability in patients with CLL and increased mutational load in B cell-derived lymphoma cells.

### Loss of Lamin B1 contributes to elevated DNA damage and recruitment of DDR proteins

The observed association between Lamin B1 and GI in CLL samples led us to investigate the effect of Lamin B1 reduction on GI and its impact on recruiting DNA repair proteins in GC-B cell-derived lymphoma cell lines and *in vivo* during B cell development in GCs.

We first generated doxycycline-inducible Lamin B1 knockdown in GC-derived lymphoma cell lines. Transduced BL2 (Burkitt’s lymphoma) and OCI-LY8 (GCB DLBCL) cells were analysed for GFP positivity by flow cytometry upon addition of doxycycline (DOX) (Fig. S2A-B), and the efficacy of short-hairpin RNA (shRNA)-mediated Lamin B1 knockdown was assessed by western blot analysis (Fig. S2C). Alkaline single-cell electrophoresis (comet) assay provided an overview of DNA damage after Lamin B1 knockdown in BL2 and OCI-LY8 lines. OCI-LY8(Singh, McCoy et al., 1988). We observed that Lamin B1 reduction led to higher tail moments in shLMNB1 cells compared to control, revealing an overall increased DNA damage (Fig. 2A-B).

**Figure 2.**
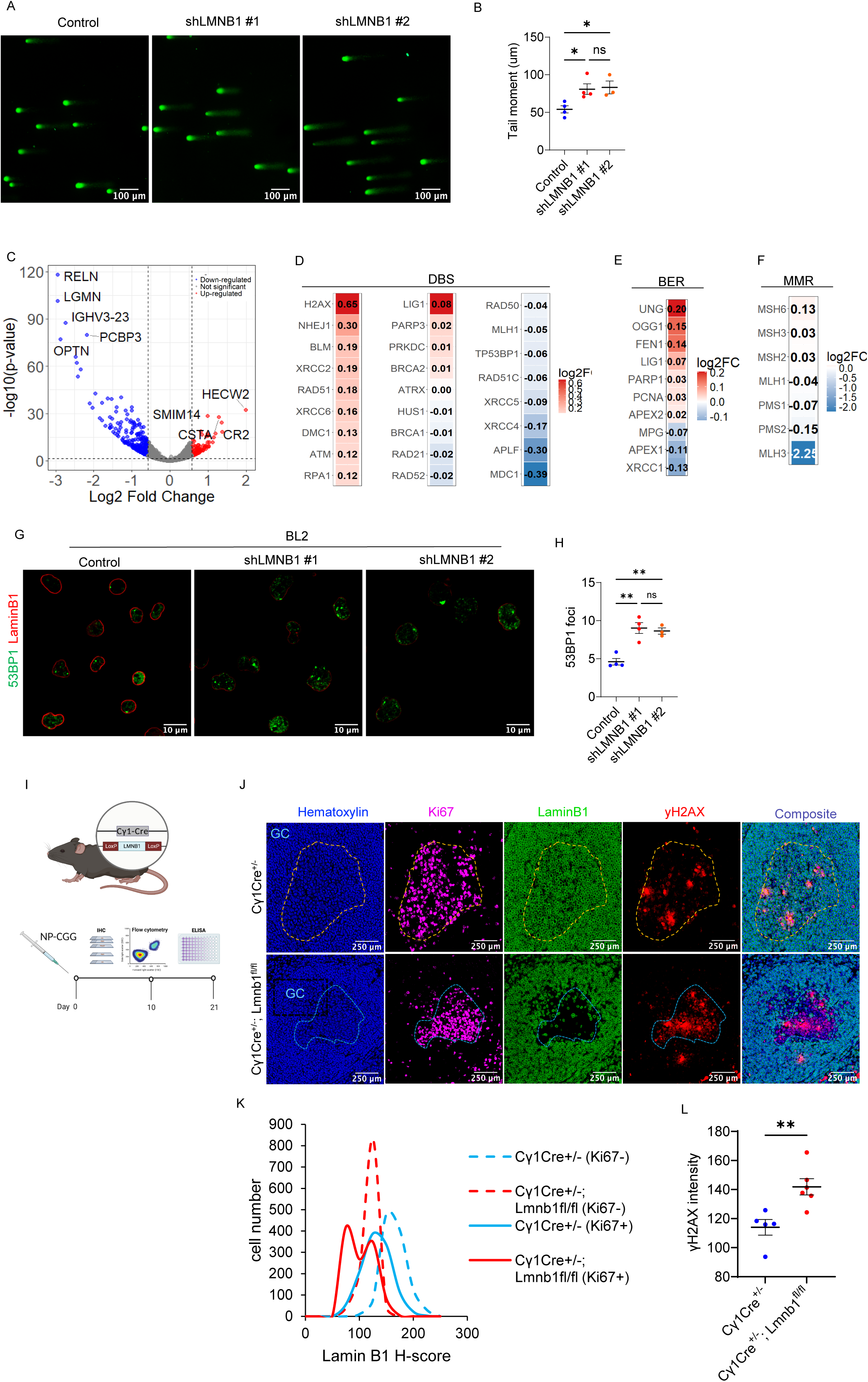
Lamin B1 reduction translates into increased GI. **(A)** Representative COMET assay images of BL2 control cells and stable shRNA Lamin B1 sequences. (**B**) Quantification of COMET tail length. Data represent three independent experiments. (**C**) Volcano plot comparing gene expression in shLMNB1 versus control BL2 cells. Significantly upregulated genes (padj < 0.05, log₂FC > 0.5) are shown in red; significantly downregulated genes (padj < 0.05, log₂FC < -0.5) in blue. (**D-F**) Log₂ fold change (shLMNB1 vs control) of genes involved in: BER (**D**), DSB (**E**), MMR (**F**) as introduced in Figure 1A-B and Supplementary Figure 1A. (**G**) Immunofluorescence images showing Lamin B1 (red) and 53BP1 (green) staining in control and shLMNB1 cells. (**H**) Histogram showing 53BP1 distribution from panel g. *n* = 3 independent experiments. (**I**) Schematic of the conditional Lamin B1 knockout mouse model in germinal centre (GC) B cells and associated analyses. (**J**) Representative image of multiplex immunohistochemistry showing the Lamin B1 reduction in germinal centres (Ki67+ structures) upon control (*Cγ1Cre⁺/⁻*) and Lamin B1 floxed (*Cγ1Cre⁺/⁻; Lmnb1^fl/fl^*) mice in Ki67⁻ (non-GC) and Ki67⁺ (GC) regions. (**L**) Quantification of γH2AX intensity in GC regions of control vs Lamin B1 knock-out mice.

GI and transcriptional activity are tightly interconnected (Milano, Gautam et al., 2024). Previous studies have demonstrated that Lamin B1 contributes to gene repression within LADs, and that removing Lamin B1 modulates the transcriptional profile in various cancer models (Jia, Vong et al., 2019, Reddy et al., 2008, Reilly et al., 2022). To gain insight into the transcriptional landscape modulated by Lamin B1 in our model, we performed RNA-seq analysis of shLMNB1-treated BL2 cells (Fig. S2D). Using the DESeq2 algorithm, we detected differentially expressed genes in BL2 cells upon Lamin B1 silencing. 526 genes were significantly dysregulated (padj <0.05, Log2FC ±> 0.5) (Fig. 2C; Table S1). We examined changes in the expression of BER, DSB, and MMR pathway genes by comparing their log₂ fold change between shLMNB1 and control conditions. Overall, the results reveal a significant upregulation of genes involved in the DSB and BER pathways, indicating that the loss of Lamin B1 activates this DNA repair mechanism (Fig. 2D-F). As an additional control, we assessed Lamin B1 protein levels before performing the RNA-seq experiment (Fig. S2D) and further validated the expression of LMNA, LMNB1, LMNB2, and APOBEC family genes in our RNA-seq dataset (Fig. S2E, F). The results showed that the relative expression of all queried genes remained unchanged in shLMNB1 compared with control BL2 cells, except for Lamin B1, whose expression was significantly reduced under the knockdown condition. These findings indicate that the changes observed in our system are specific to Lamin B1 depletion.

To validate the upregulation of key proteins from the DSB pathway, we performed immunofluorescence and western blot analysis of γ-H2AX and 53BP1. A marked increase in 53BP1 foci per cell was observed following Lamin B1 depletion (Fig. 2G, H; Fig. S2G), indicating an enhanced DNA damage response in BL2 and OCI-LY8 cell lines. Similarly, phospho-γ-H2AX (Fig. S2H) levels were elevated in OCI-LY8 Lamin B1 knockout cells, further supporting the activation of DNA repair pathways.

Next, we generated a GC B cell-specific *LMNB1* knockout mouse model to evaluate Lamin B1’s impact on DSBs under physiological conditions (Fig. 2I). Using multiplex immunohistochemistry (mIHC) (Fig. 2J), we identified GCs histologically using the Ki67 marker. We then evaluated the levels of Lamin B1 (Fig. 2K) and phospho-γ-H2AX (Fig. 2L). Our observations showed that the depletion of Lamin B1 alone is sufficient to increase the γ-H2AX signal. Additionally, we isolated PNA+ (GC B) and PNA-(non-GC B) cells (Fig. S2I) and confirmed by immunofluorescence that γ-H2AX was increased in Lamin B1 KO PNA+ cells, supporting our IHC findings. Altogether, these results suggest that Lamin B1 plays a crucial role in maintaining genomic integrity, and its loss alone leads to an increase in DNA double-strand breaks or replication stress.

### Genome-wide profiling of DSB topology in human and mouse Lamin B1-depleted B cells

By modulating chromatin accessibility, Lamin B1 helps orchestrate GC B cell programmes and preserves GC architecture (Sallan, Filipsky et al., 2025). Alterations in genome accessibility and transcriptional programmes often manifest as phenotypic changes at the tissue level. To determine whether these molecular alterations were reflected in GC organisation, we examined spleen architecture by immunofluorescence staining for BCL6, which delineates germinal centres (Fig. 3A). We observed that *Cγ1Cre^+/-^;Lmnb1^fl/fl^* mice displayed a higher number of GCs compared with *Cγ1Cre^+/-^* control (Fig. 3B), although the average GC size, measured by area, was significantly reduced (Fig. 3C).

**Figure 3.**
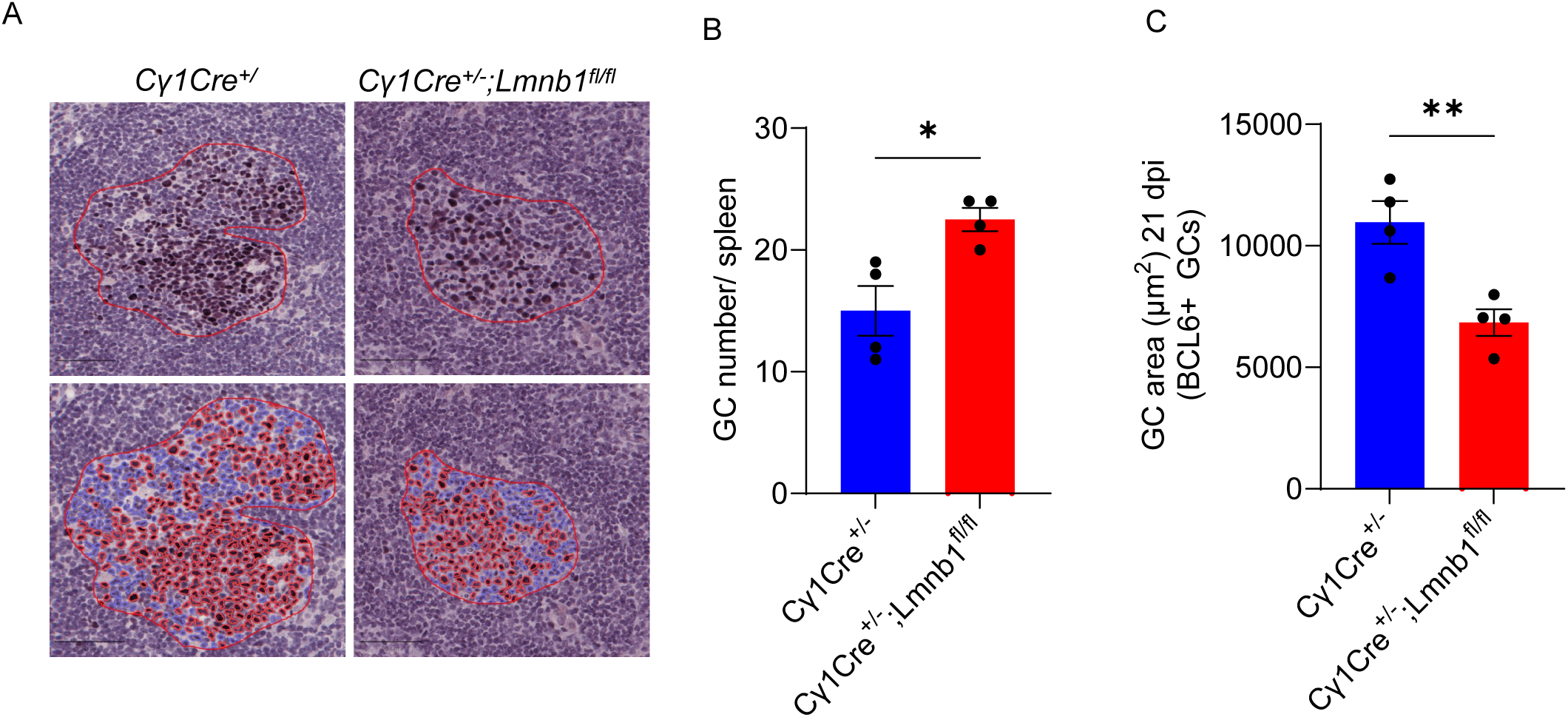
Lamin B1 depletion in GC B cells results in GC phenotypic changes. (**A**) Representative images of BCL6 immunohistochemistry on *Cγ1Cre^+/-^* and *Cγ1Cre^+/-^ ;Lmnb1^fl/fl^* spleens 21 days post immunisation. Cell detections (Red = positive BCL6 staining; blue = negative BCL6 staining) in GCs. (**B**) Barplot showing GC number measured by BCL6+ clusters (>10 cells) per spleen section. Comparison of *Cγ1Cre^+/-^* (n=4 mice) and *Cγ1Cre^+/-^ ;Lmnb1^fl/fl^* (n=4 mice) in at least two independent immunisations. Two-tailed t-test (p-value <0.01). (**C**) Barplot showing GC area measured using BCL6+ clusters (>10 cells) per spleen section. Comparison *Cy1Cre^+/-^;Lmnb1^fl/fl^* (n=4 mice) and *Cγ1Cre^+/-^* (n=4 mice) in at least two independent immunisations. Two-tailed t-test (p-value <0.01).

Given the architectural and transcriptional disruption observed in *Cγ1Cre^+/-^;Lmnb1^fl/fl^* GCs, we next sought to determine whether these changes were accompanied by increased genomic instability. We performed genome-wide mapping of DNA DSBs to identify which Lamin B1-protected regions were preferentially prone to breakage, revealing structural vulnerabilities. For this, we employed Breaks Labelling in Situ and Sequencing (BLISS) (Fig. 4A; Fig. S3A, B) to map regions of increased DNA fragility (Bouwman, Agostini et al., 2020).

**Figure 4.**
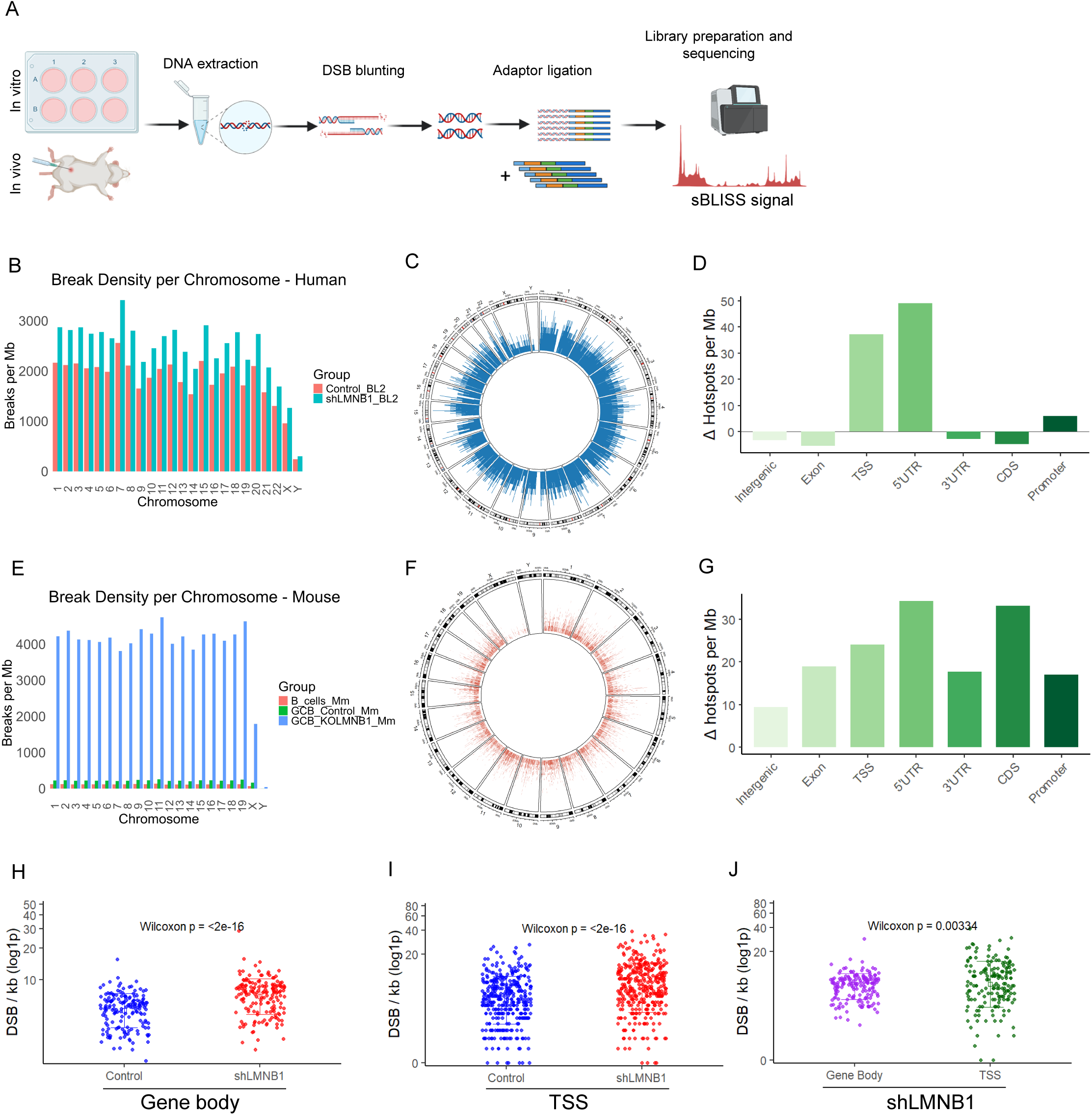
Lamin B1 protects the genome from high-density DNA breaks. **(A)** Schematic diagram illustrating the experimental workflow for generating sBLISS data. (**B**) Bar plot showing the distribution of DSB density (breaks per megabase) across chromosomes in BL2 cells under control and shLMNB1 conditions. (C) Circoplot showing the unique hotspot distribution in shLMNB1 compared to the control in BL2 cells. (**D**) Barplot showing increase/decrease in hotspots in shLMNB1 compared to control in seven genomic features. (**E**) Bar plot showing DSB density (breaks per megabase) across chromosomes in *ex vivo* samples: non-germinal centre Naïve B cells (B_cells_Mm), *Cγ1Cre^+/-^* GC B cells (GCB_Control_Mm), and *Cγ1Cre^+/-^; Lmnb1^fl/fl^* GC B cells (GCB_KOLMNB1_Mm). (**F**) DSB unique hotspots distribution in *Cγ1Cre^+/-^;* Lmnb1^fl/fl^ compared to *Cγ1Cre^+/-^; Lmnb1^fl/fl^* GC B cells. (**G**) Bar plot showing increases/decreases in hotspots in shLMNB1 compared to control across seven genomic features. (**H**) Dotplot showing DSB/Kb in DEGs body regions comparing control and shLMNB1. (**I**) Dotplot showing DSB/Kb in DEGs TSSs comparing control and shLMNB1 (**J**) Dotplot comparing DSB/Kb in gene body and TSS in shLMNB1 condition using DEGs from RNAseq .

The BL2 cell line, mouse splenic germinal centre B cells (B220⁺, CD95⁺, GL7⁺) and naïve B cells (B220⁺, CD95⁻, GL7⁻) were used for the analysis. Principal Component Analysis (PCA) revealed replicate consistency and sample variability between conditions in *in vivo* and *in vitro* scenarios (Fig. S3C, D).

We first quantified DSBs per megabase (Mb) across individual chromosomes. Our *in vitro* BL2 Lamin B1 knockout model showed increased DSBs across all chromosomes (Fig. 4B) . Regional analysis revealed that Lamin B1 deficiency led to a broader increase in DSB burden, particularly at 5’ and 3’UTR, TSS, and exons (Fig. S3E). When analysing DSB hotspots, we observed that Lamin B1 depletion *in vitro* led to a markedly higher number of unique genome-wide DSB hotspots compared withcontrol cells (Fig. 4C). Complementary to this, the distribution of differential hotspots against the genomic background (control BL2) *in vitro* appeared non-random, being preferentially concentrated around genes (TSS, 5’UTR and promoter regions (Fig. 4D). Consistent with the *in vitro* model, the *in vivo* Lamin B1 knockout exhibited a similar, widespread accumulation of DSBs, indicating a global increase in genomic fragility (Fig. S3F).

In mouse samples, the break density per chromosome and DNA region, as well as the number of DSB hotspots, were considerably higher in *Cγ1Cre^+/^-;Lmnb1^fl/fl^* GC B cells compared with control GC B cells (Fig. 4, E–F). This suggests that Lamin B1 loss has a more profound effect on DSB susceptibility in non-malignant GC B cells. We next tested the distribution of unique DSB hotspots in our Lamin B1 knock-out GC B cells against the *Cγ1Cre^+/-^* background. Notably, the results revealed an accumulation of DSB hotspots across all genomic features analysed. The most significant enrichment was observed at transcription start sites (TSS), 5′ untranslated regions (5′UTRs), and coding sequences (CDS), confirming that DNA breaks arising from Lamin B1 loss are non-randomly distributed across the genome (Fig. S3F).

Upon LMNB1 loss, hotspot signal values rise in both the BL2 cell line and mouse GC B cells. The genomic distribution of hotspots, together with the dynamic range of their break frequencies, is largely conserved in human and mouse samples (Fig. S3G). This cross-species concordance indicates a shared breakage-prone landscape that becomes unmasked when Lamin B1 function is disrupted.

We next leveraged our BL2 RNA-seq data to assess DSB distribution across gene bodies and transcription start sites (TSSs) of DEGs. Upon *LMNB1* reduction, DSB accumulation increased at both TSSs and gene bodies of DEGs (Fig. 4H, I). Furthermore, within the LMNB1 knockdown condition, TSSs displayed a significantly higher DSB density per kilobase than gene bodies (Fig. 4J), indicating that DSB formation is not random and is preferentially enriched at promoter-proximal regions, consistent with the sBLISS findings shown in Fig. 4D.

Finally, we asked whether genes commonly mutated in DLBCL (Coyle, Dreval et al., 2025) coincide with DSBs revealed in our system. Out of 142 DLBCL genes, 134 (≈94%) overlapped with at least one DSB, and 76 (≈54%) harboured hotspots specific to the shLMNB1 condition (Table S2). These findings are consistent with a model in which a subset of mutations arises from impaired DSB repair associated with LMNB1 downregulation.

Together, these data indicate that Lamin B1 safeguards genome stability in germinal centre B cells by limiting the formation of non-random, promoter-proximal DNA DSBs. Thus, Lamin B1 functions as a key structural determinant of genome integrity in B cells.

### Lamin B1 loss enhances DNA fragility at LADs and rewires functional gene networks

We next sought to investigate how the reduction of Lamin B1 impacts the functional genomic landscape in lymphoma and in GC B cells. We utilised the sBLISS data to examine the genomic distribution of unique DSB hotspots in Lamin B1-depletedt cells, both *in vitro* (Fig. 5A) and *in vivo* (Fig. 5B), by dichotomising the genome into LAD and non-LAD regions. In *in vitro* settings, we found that 80.7% of DSB hotspots occurred within LADs, divided between 39.3% in constitutive LADs (cLADs) and 41.4% in facultative LADs (fLADs). In comparison, only 19.3% of hotspots were localised outside LADs. In contrast, the distribution of DSB hotspots in mouse GC B cells exhibited a markedly different pattern, with only 33.6% of hotspots located within LADs. It is important to note, however, that in mice the distinction cLADs and fLADs is less clearly defined, and a subset of fLADs may therefore be annotated within the non-LAD category.

**Figure 5.**
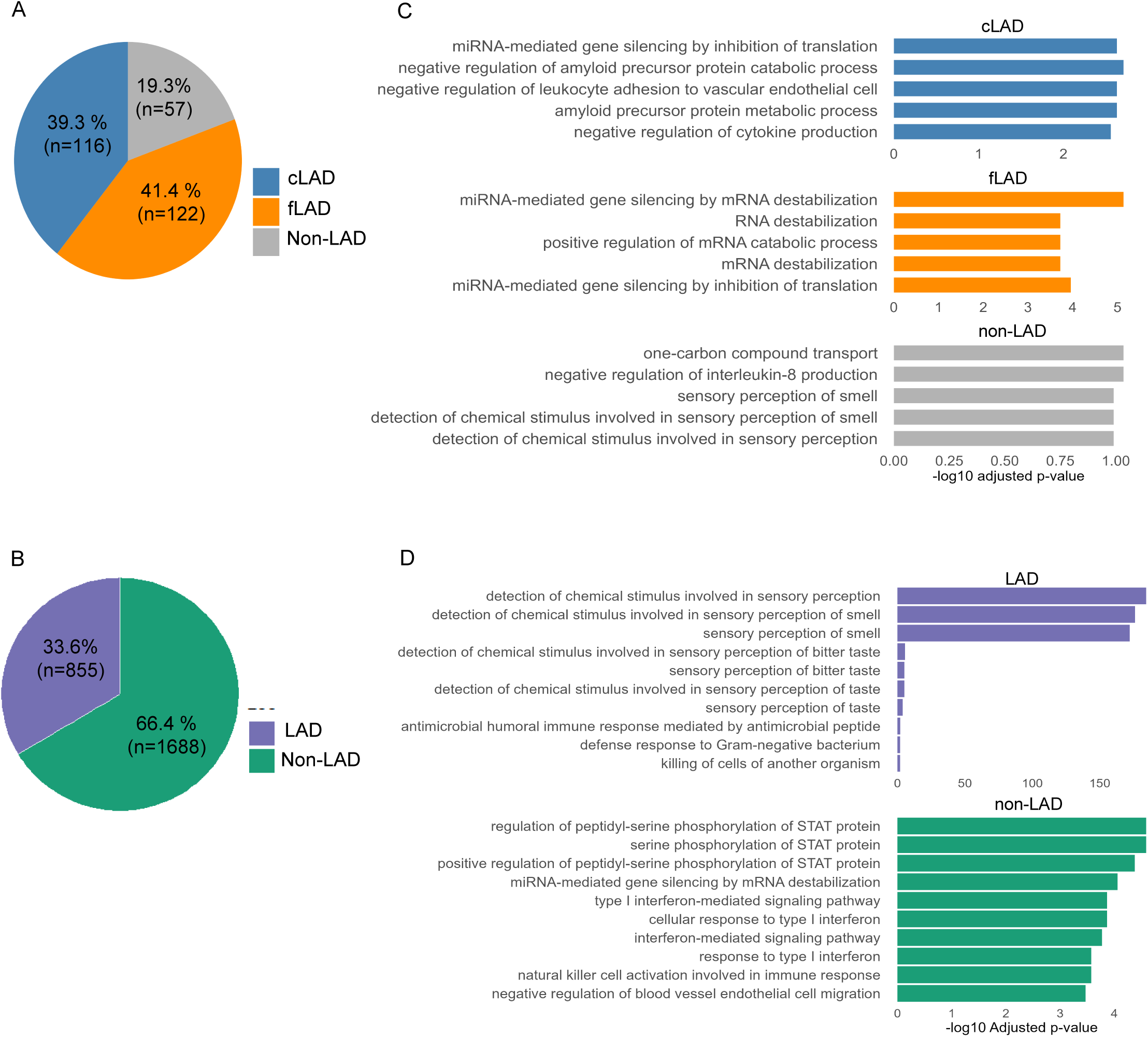
Lamin B1 downregulation impacts genome architecture rewiring the functional genomic landscape of the cell. (**A**)Pie chart showing the genomic distribution of unique Lamin B1–dependent DSB hotspots classified into LADs (constitutive LAD -cLAD- and facultative LAD -fLAD-) and non-LAD regions in BL2 cell line. (**B**) Pie chart showing the genomic distribution of unique Lamin B1-dependent DSB hotspots classified into LADs and non-LAD regions in *Cγ1Cre^+/-^;Lmnb1^fl/fl^*GC B cells. (**C**) Gene Ontology (GO) Biological Process (BP) enrichment of genes associated with cLAD, fLAD and non-LAD from A. (**D**) Barplot quantification of significantly enriched GO terms from genes associated with LAD vs non-LAD contexts from B.

To gain deeper insight into the biological processes potentially affected by DSBs, we performed Gene Ontology (GO) enrichment analysis on genes associated with DSB hotspots located within LADs and non-LADs (Fig. 5C). Genes that were both located within cLADs and affected by DSB hotspots were predominantly enriched in processes related to immune regulation and protein metabolism. By contrast, genes within facultative fLADs were strongly associated with RNA regulatory processes, including mRNA destabilisation and catabolism. This suggests that developmentally regulated regions undergoing chromatin remodelling may be particularly susceptible to DSBs that impact RNA turnover. Non-LAD-associated genes with DSBs exhibited overall weaker enrichment. Their top GO terms were related to chemical stimulus detection and sensory perception, indicating a distinct and potentially more stochastic damage pattern in non-LAD-associated regions.

The landscape in mouse GC B cells presented a different pattern. Genes located within LADs and affected by DSBs were predominantly enriched in sensory-related processes. In contrast, DSB-associated genes in non-LAD regions showed strong enrichment for immune-related pathways, including STAT protein phosphorylation, type I interferon signalling, natural killer (NK) cell activation, and immune cell migration (Fig. 5D). These findings suggest that while LADs in mice tend to harbour DSB-prone loci linked to sensory functions, genomic instability outside the lamina preferentially affects immune regulatory networks.

### Lamin B1 loss promotes DSBs at AICDA and novel intergenic motifs

GC B cells are the origin of many lymphomas and rely on AID as the primary mutator enzyme (Teater, Dominguez et al., 2018). We therefore investigated how Lamin B1 depletion affects DNA damage at AID targets and newly *de novo* motifs. We first focused on AID hotspots (WRC, WRCY, and WGCW) (Kohli, Maul et al., 2010) using DSB hotspots from the *Cγ1Cre⁺/⁻;LMNB1^fl/fl^*mouse model. Using the CentriMo algorithm, we determined the location probability of these motifs within the DNA sequence associated with DSBs (Fig. 6A; Fig. S4A). As expected, the AID target motif sequence was located near the centre of the sequence, corresponding to the original DSB. A non-AID motif sequence (GAGA) was not located at the centre of the DSB sequence, confirming the specificity of the AID-induced DSB site in GC B cells.

**Figure 6.**
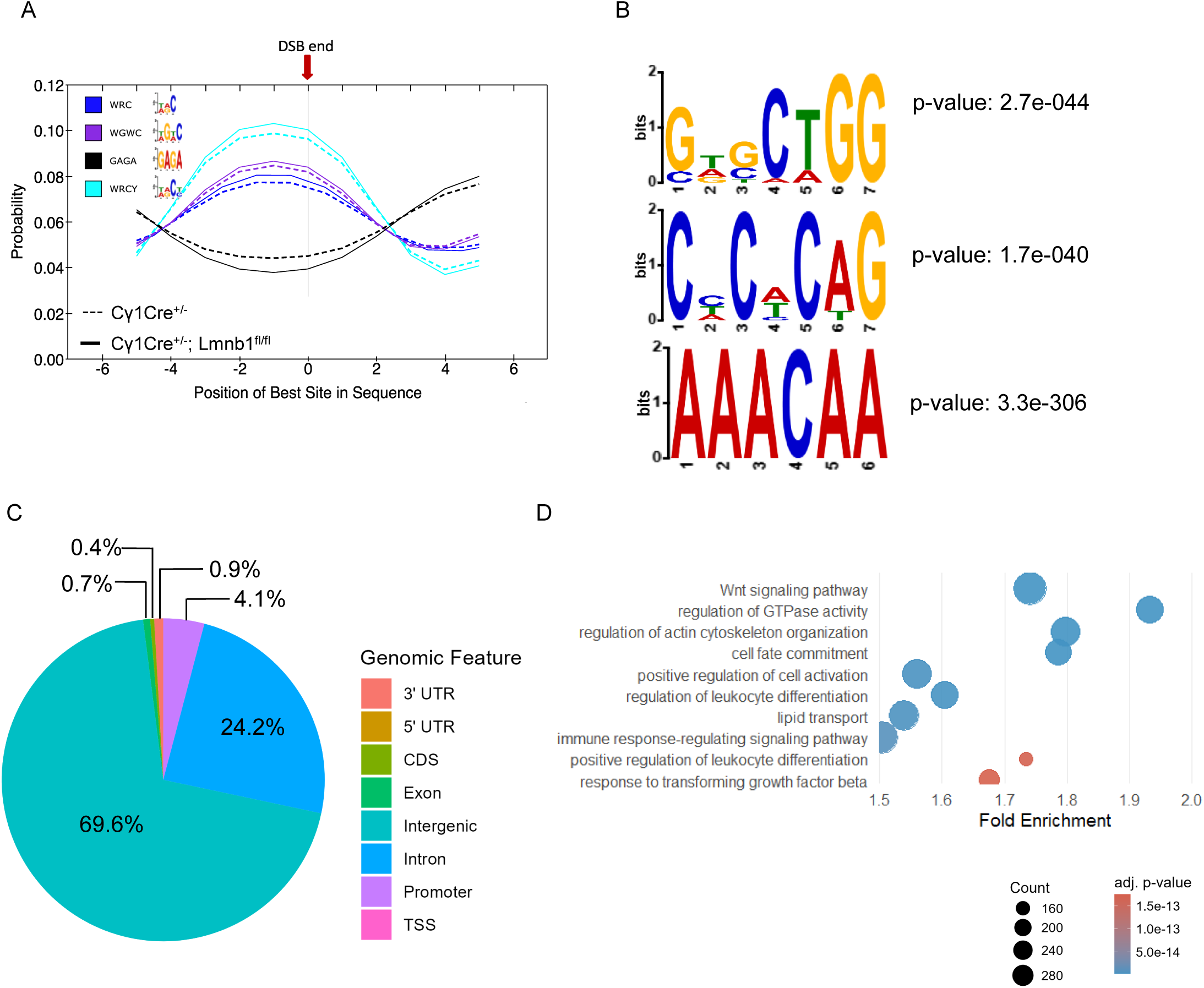
Lamin B1 loss promotes DSBs at AICDA motifs and defines *de novo* nucleotide patterns. (**A**) AID hotspots (WRC, WRCY, and WGCW) location probability associated with DSBs in *Cγ1Cre^+/-^; Lmnb1^fl/fl^*compared to *Cγ1cre^+/-^* GC B cells. (**B**) Top 3 enriched *de novo* motifs in *Cγ1cre^+/-^; Lmnb1^fl/fl^* GC B cells. (**C**) Pie chart showing percentage of *de novo* motifs found at different genomic regions in *Cγ1cre^+/-^; Lmnb1^fl/fl^*. (**D**) GO enrichment results showing deregulated biological processes associated with genes annotated near motif locations (±3 kb of a TSS) derived from the discovered motif sites.

Using MEME Suite (v5.5.8), we identified several *de novo* motifs significantly enriched around Lamin B1-associated DSB hotspots (Fig. 6B; Fig. S4B). The top motifs included GWGCTGG (E = 1.7e-40), found at 2,710 sites and corresponding to the canonical SPI1/PU.1 and MEF2C binding sites, and CHCWCAG (E = 1.7e-40), a GC-rich motif frequently observed in enhancer-like regions. Additionally AAACAA and TTTCTT, which are AT-rich motifs that may reflect fragile genomic regions or regulatory repressor elements. Notably, the GWGCTGG motif contains a central CTGG core that was highly conserved and corresponds to known haematopoietichaematopoietichaematopoietic transcription factor (TF) binding motifs.

To determine their genomic distribution, we mapped the *de novo* motifs and found that most reside in intergenic (69.6%), intronic (24.2%), and promoter (4.1%) sequences (Fig. 6C). This pattern indicates that DSBs arising after Lamin B1 depletion occur non-randomly and are enriched in regulatory regions, underscoring the nuclear lamina’s role in safeguarding genome integrity.

To further extend the analysis, the *de novo* motifs were then matched to known TF motifs using Tomtom (MEME Suite). To contextualise the biological relevance of matched TFs, we utilised the CistromeDB, querying publicly available ChIP-seq data from different cell types to evaluate TF binding site enrichment near our input regions. GIGGLE similarity scores were computed. High-scoring factors, such as LEO1, MED1, KMT2B, and STAT5, emerged as candidates potentially involved in DSB-associated regulatory processes (Fig. S4C). To understand the chromatin context of these breaks, we applied GIGGLE enrichment analysis against reference epigenomic datasets again.

Our analysis showed that DSB hotspots unique to Lamin B1-deficient GC B cells were strongly enriched in regions bearing heterochromatin-associated histone marks, particularly H3K9me3 and H3K27me3 (Fig. S4D). By contrast, we observed only moderate overlap with core histone H3 and enhancer-associated H3K27ac, and minimal enrichment for transcription-elongation marks (H3K36me3, H3K79me2) or less common histone variants (Fig. S4D). Together, these patterns indicate that Lamin B1 loss biases DSB formation towards repressive chromatin environments in B cells.

To identify biological processes that are potentially affected by motif-associated DNA damage, we extended the genomic coordinates of *de novo* discovered motifs by ±3 kb and mapped them to the nearest TSS. This mapping yielded a set of genes near the motif hotspots. Gene Ontology enrichment analysis revealed that these genes were significantly associated with pathways involved in actin cytoskeleton organisation, cell fate commitment, and Wnt signalling (Fig. 6D; Table S3): processes implicated in haematological tumour progression and plasticity (Beguelin, Popovic et al., 2013, Gutierrez, Tschumper et al., 2010, Velazquez-Avila, Balandran et al., 2019). Additionally, enrichment of immune-related terms, such as leukocyte differentiation and immune response–regulating signalling, further suggests that DSB accumulation at motif-associated regions can shape both structural and immunomodulatory programmes in lymphoma.

### Decreased *LMNB1* expression is associated with poor clinical outcomes of DLBCL

In haematological malignancies, including CLL and acute myeloid leukaemia (AML), reduced *LMNB1* expression is associated with adverse clinical prognosis (Klymenko et al., 2018, Reilly et al., 2022). Considering the profound involvement of *LMNB1* in processes implicated in haematological tumour progression and plasticity (Fig. 6D), we next investigated the clinical significance of Lamin B1 in GC lymphomas, including DLBCL (Schmitz, Wright et al., 2018) (Fig. 7A). Remarkably, we observed a significantly shorter 5-year survival rate in all patients with low *LMNB1* expression, as stratified by the median (Fig. 7B). We next investigated whether the *LMNB1*-mediated survival difference is associated with the cell of origin and genetic subtype.

**Figure 7.**
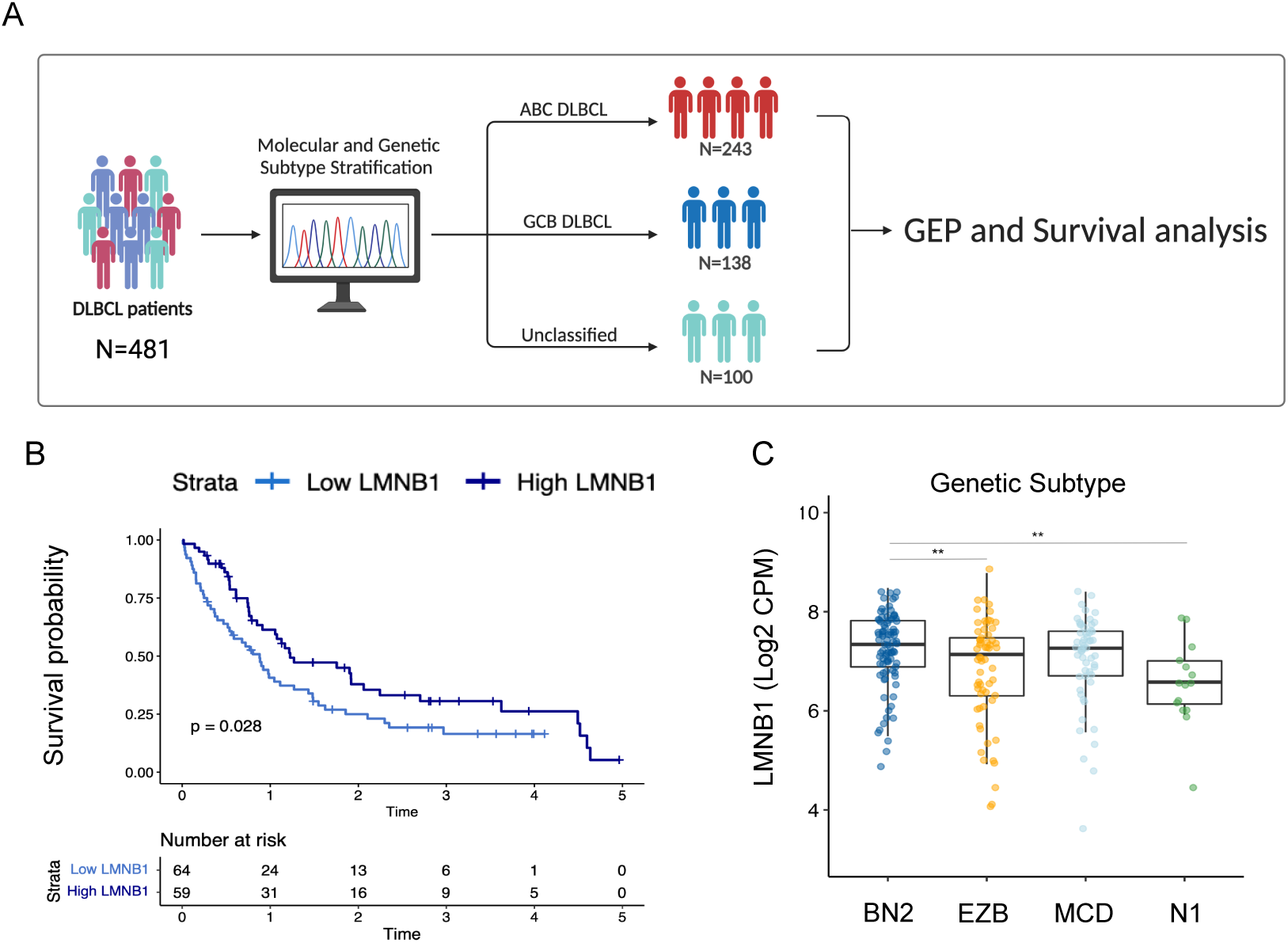
Decreased *LMNB1* expression is associated with worse clinical outcome in DLBCL. (**A**) Workflow diagram of clinicopathological analysis in DLBCL patients. (**B**) Kaplan-Meier curve estimates of 5-year PFS based on median LMNB1 expression in DLBCL patients (N=123). (**D**) *LMNB1* expression is decreased in N1 and EZB genetic DLBCL subtypes (N=220). Unpaired t-test was used for the statistical analysis \**p* ≤ 0.05; \*\**p* ≤ 0.01; ns - not significant.

While *LMNB1* expression did not vary across stages in the ABC subtype (Fig. S5A), a stage-associated decline was observed in GCB DLBCL, reaching its lowest levels at intermediate stages (II–III) before stabilising at stage IV (Fig. S5B). This pattern suggests that *LMNB1* loss may occur early during disease progression and is not further exacerbated at the advanced stage. Genetic classification revealed that *LMNB1* expression was reduced in the N1 and EZB subtypes (Fig. 7C), consistent with the notion that diminished *LMNB1* contributes to a more malignant phenotype. Taken together, these findings indicate that low *LMNB1* expression is associated with poor prognosis in DLBCL and is functionally linked to the GCB and the N1 and EZB genotypes.

## Discussion

While controlled DNA damage is essential for humoral immunity, it can inadvertently generate chromosomal translocations and other genomic alterations that initiate lymphomagenesis (Basso & Dalla-Favera, 2015, Daniel & Nussenzweig, 2013). The mechanisms safeguarding against such genomic instability remain incompletely understood. Here, we investigated the role of Lamin B1 in maintaining genome stability in human lymphoma cells and normal murine GC B cells. We found that Lamin B1 depletion in B cells was associated with widespread transcriptional reprogramming, clustering of DNA double-strand breaks (DSBs), and poor survival outcomes in both CLL and DLBCL.

Low *LMNB1* expression and GI were independently reported as adverse prognostic factors in the CLL8 and REACH CLL trial cohorts (Bloehdorn et al., 2021, Klymenko et al., 2018). Our findings showed that *LMNB1* expression is particularly reduced in CLL cases of the GI subtype (REACH trial cohort), providing evidence that loss of Lamin B1 is associated with genomic instability in CLL and possibly other B-cell-derived malignancies.

Upon B cell activation, reduced Lamin B1 nuclear incorporation was shown to be accompanied by a higher mutational load within the *IghV* cluster (Klymenko et al., 2018). In addition, mutational clustering of primarily C to T substitutions, consistent with the AID mutagenic signature, occurs outside the IghV region in Lamin B1-depleted lymphoma cells. However, similar mutational patterns in BL2 *AID^wt^* and BL2 *AID^-/-^* cells suggest the potential involvement of other APOBEC deaminases such as *APOBEC3C* and *APOBEC3G*, which are also known drivers of mutagenesis in myeloma (Talluri, Samur et al., 2021).

Lamin B1 depletion from the nuclear periphery is associated with changes in chromatin organisation, DNA accessibility, and damage dynamics (Chapman, Filipsky et al., 2023, Noguchi, Ito et al., 2021, Pascual-Reguant et al., 2018, Shah et al., 2013). Here, we also show that increased accessibility in Lamin B1-deficient cells drives a targeted, AID-independent rise in mutations outside Ig loci, supported by the non-uniform distribution of SNV clusters. To our knowledge, this is the first demonstration that reduced *LMNB1* expression elicits spontaneous DNA damage in B cells *in vitro* and *in vivo*.

The increased localisation of 53BP1 in response to Lamin B1 depletion is also in agreement with the report that Lamin B1 controls the recruitment of 53BP1 to the damaged site in SV40-transformed human fibroblasts and U2OS cells (Etourneaud et al., 2021). Interestingly, the NHEJ pathway is a primary repair mechanism in B cells and is biased towards DSBs located within relaxed chromatin (Schep, Brinkman et al., 2021). Therefore, our data suggest that the fast and error-prone NHEJ pathway is involved in the repair of DSBs due to increased 53BP1 localisation in Lamin B1-depleted cells.

Next, Lamin B1 is essential for mouse development and cell differentiation, as established in several *in vivo* models (Chen, Yang et al., 2019, Jia et al., 2019, Vergnes, Peterfy et al., 2004). To investigate its role in GC biology while avoiding the deleterious effects of global *LMNB1* deletion, we generated a conditional GC B cell-specific *LMNB1* knockout mouse model.

Our model demonstrates that Lamin B1 reduction alone is sufficient to induce a widespread increase in DNA DSBs in GC B cells. Furthermore, by delineating the genomic landscape of this damage, we revealed that DSB hotspots accumulated predominantly within transcriptionally active and accessible chromatin regions, particularly near promoters.

Notably, accessible genes in control GC B cells did not exhibit DSBs at the stringent detection thresholds applied, indicating a tightly regulated chromatin architecture that normally restrains break formation. In contrast, 42.8% of the genes that became accessible upon Lamin B1 depletion coincided with DSBs, underscoring a breakdown in genome surveillance following LMNB1 loss. Strikingly, integration with a high-resolution human DSB map revealed that 96% (134/142) of the genes most frequently mutated in DLBCL contained at least one DSB, and 53% (76/142) harboured hotspots unique to the LMNB1-depleted condition. These patterns suggest a protective functional role for Lamin B1 at key lymphoma driver loci.

As expected, stratification of DSBs by lamina association revealed that *in vitro* Lamin B1 depletion predominantly induced DSBs within LAD-associated genes, which, in turn, showed transcriptional deregulation. While Lamin B1 is known to regulate genome positioning in both LADs and non-LADs (Shah et al., 2013, Zheng, Hu et al., 2018), our data show that its loss also increases DSBs in non-LAD regions, highlighting its role as a broader chromatin protector beyond the classical LADs.

Interestingly, DSBs in *LMNB1*-deficient GC B cells were primarily confined to non-LAD regions under physiological conditions. This pattern may arise, at least in part, from the biology of GC B cells, in which Lamin B1 expression naturally declines as antigen-experienced B cells enter the germinal centre and undergo somatic hypermutation. During this transition, LADs are likely remodelled, exposing non-LAD regions to greater fragility upon complete Lamin B1 loss. An additional consideration is that LADs are less precisely defined in mouse compared with human genomes, which may contribute to the apparent enrichment of DSBs outside LAD domains.

We found that DSBs were preferentially enriched around transcription start sites (TSSs) in Lamin B1-deficient cells. This distribution aligns with the well-characterised off-target activity of AID outside immunoglobulin loci (Bransteitter, Pham et al., 2004, Goyenechea, Klix et al., 1997, Pham, Bransteitter et al., 2003) which targets highly transcribed genomic regions with specific sequence preferences.

Notably, Lamin B1 loss *in vivo* also led to the emergence of *de novo* DNA motifs enriched at DSB sites. These motifs corresponded to transcription factors involved in haematopoiesis and heterochromatin regulation, indicating that Lamin B1-dependent chromatin architecture may limit DNA fragility by restricting transcription factor access to vulnerable genomic regions.

Finally, extending our mechanistic insights to patient data, we found that *LMNB1* expression is diminished in the N1 and EZB genetic subtypes and declines progressively with advancing clinical stage within the GCB DLBCL subtype. Consistent with its role in safeguarding genome stability, reduced *LMNB1* expression was associated with inferior clinical outcomes, functioning as a negative prognostic marker whereby low expression predicted shorter PFS across DLBCL genetic subtypes.

## Materials and Methods

### Cell lines and Lamin B1 knockdown

Burkitt’s lymphoma BL2, GCB-DLBCL OCI-LY8, and human embryonic kidney HEK293 cells were obtained from the German Collection of Microorganisms and Cell Culture (Deutsche Sammlung von Mikroorganismen und Zellkulturen, DSMZ). BL2 and OCI-LY8 were cultured in RPMI 1640 medium supplemented with L-Glutamine, 10% FBS and 1% penicillin/streptomycin (PS). HEK293 cells were maintained in Dulbecco’s Modified Eagle’s medium supplemented with 10% FBS and 1% PS. The cell lines were maintained at 37°C with 5% CO_2_, passaged every 48 h, and routinely screened for mycoplasma contamination. To generate stable shRNA Lamin B1 knockdown cell lines, BL2 cells were transduced with lentiviral particles containing specific shRNA sequences (shLMNB1 #1: [V3SH11252-224783461], shLMNB1 #2: [V3IHSMCG_8167334]). For validation, OCI-LY8cells were transduced with shRNA (shLMNB1 #1: [V3SH11252-224783461] against LMNB1 mRNA using a calcium phosphate transfection protocol. Briefly, HEK293 cells were transfected using a Dharmacon trans-lentiviral shRNA packaging kit with calcium phosphate transfection reagents (TLP5913), according to the manufacturer’s protocol. BL2 or OCI-LY8cells were transduced with lentivirus-containing supernatant and positively selected with puromycin at a 2.5 µg/mL concentration for 72 h. Target cell lines containing lentiviral vectors were incubated with 500 ng/mL of doxycycline (DOX) to induce shRNA expression. The percentage of GFP-expressing cells was assessed using flow cytometry 24 h after the addition of DOX, and Lamin B1 knockdown was assessed 72 h later. For transient Lamin B1 knockdown, cells were subjected to electroporation as previously described (Klymenko et al., 2018). Lamin B1 nuclear incorporation and total protein levels were assessed using IF and western blotting, respectively. Cell counts and viability were measured by acridine orange/propidium iodide staining using a Luna dual fluorescent cell counter (Logos Biosystems).

### Western Blotting

Total cell lysates were prepared by lysing harvested cells in 2X NuPAGE LDS sample buffer with protease and phosphatase inhibitors. Lysates were then sonicated for 15 cycles (30 s on/off) using a Bioruptor Pico (B01060010). Denatured proteins were separated by electrophoresis and transferred onto methanol-activated PVDF membranes. The membrane was then blocked with 3% BSA/TBS-T for one hour and incubated with the primary antibody at the specified dilution and duration (see Resource Table). The membrane was washed with TBS-T and incubated for one hour with a corresponding horseradish peroxidase (HRP)-linked secondary antibody, anti-rabbit IRDye 680RD secondary antibody, or anti-mouse IRDye 800RD secondary antibody. Images were developed with Pierce ECL Substrate using the Amersham Imager 600RGB system or Li-Cor imaging system. The images were processed using Image Studio and ImageJ software.

### Single-cell electrophoresis

The single-cell electrophoresis (COMET) assay was performed according to the protocol from the Trevigen Comet Assay kit (4250-050-K) with minor modifications. Briefly, transduced or etoposide-treated cells (positive control) were collected, washed with ice-cold PBS (no Ca/Mg), and mixed with 1% low-melting-point molten agarose (37°C) in a 1:10 cell-to-agarose ratio. The cell suspension was spread evenly onto a pre-chilled CometSlide and cooled at 4°C for at least 20 min in the dark before overnight immersion in a cold lysis solution. The slides were briefly washed with PBS and incubated in a freshly prepared 4°C alkaline solution (pH>13) for an hour in the dark. Electrophoresis was carried out in a 4°C alkaline electrophoresis solution at 21 V for 20 min with a maintained electric current of 300 mA. Immediately after electrophoresis, the slides were washed with dH_2_O for 5 min and immersed in 70% EtOH for an additional 5 min. The cells were brought to a single plane by air-drying at 37°C for at least 15 min. SYBR Gold was used to stain DNA for 30 min in the dark. Slides were then air-dried in the dark, and images were taken using a Nikon Ci-L upright fluorescent microscope at 20X magnification. TriTek CometScore 2.0, an automatic comet assay software (TriTek Corp.), was used to quantify tail moment in individual cells. Comets with a tail moment > 250 were classified as apoptotic cells and were removed from quantification.

### Flow Cytometry

To assess GFP expression, live cells were washed with PBS and incubated with DAPI (1 μg/mL) prior to flow cytometry analysis. For γ-H2AX-ser139 labelling, 1×10^6^ cells were washed with PBS (3% BSA) and fixed with ice-cold 70% ethanol (EtOH) at -20°C for 1 h. The cells were washed three times with PBS (3% BSA) and stained with anti-γ-H2AX (ser139) for 30 minutes at +4°C and 4′,6-diamidino-2-phenylindole (DAPI) (1 μg/mL) for 20 min. For the assessment of cell cycle distribution, cells were washed three times with PBS and fixed with ice-cold 70% ethanol for 1 h at -20°C. The cells were then washed, incubated with DAPI (1 μg/mL) for 20 min in the dark, and immediately analysed. Cell proliferation was assessed by FACS using Click-iT Plus EdU Flow Cytometry Alexa Fluor 488 Imaging Kit (Invitrogen/Thermo-Fisher, Paisley, UK). Logarithmically growing BL2 cells were incubated with 10 μM EdU for 2 h and processed according to the manufacturer’s instructions. The proportion of EdU+ cells was assessed using flow cytometry. At least 10,000 cells were acquired, and the data were analysed using the FlowJo software.

### WES and topology mapping

Transient Lamin B1 knockdown was performed as previously described (Klymenko et al., 2018). Whole exome sequencing (WES) with HiSeq 2 × 150 bp, with an average sequencing depth of 500X for BL2 AID^wt^, BL2 AID^-/-^, and PCL12^wt^ cells 48 h after transfection with siRNAs targeting Lamin B1. Raw reads were paired, and adapters were trimmed using fastp. Reads were mapped to the hg38 reference genome using BWA-MEM. SNVs were called with Mutect2 (v3.5), and variants found specifically in siLMNB1-treated samples were used as input for signature deconvolution using deconstructSigs. Mutational clustering was performed using KaryoplotR (v1.22.0) package.

### RNA isolation and sequencing

RNA was extracted using the RNeasy Qiagen Kit, according to the manufacturer’s protocol. BL2 cells were harvested for RNA extraction and library preparation after shLMNB1 treatment for 72 h. Paired-end sequencing was performed using Illumina HiSeq 2500 with a minimum of 30 million reads per sample. Briefly, adapter sequences were trimmed using TrimGalore (v0.6.3) and aligned to hg19 using HISAT2 (v2.0). QualiMap (v2.2.2) was used to assess the quality of alignment, and raw read counts were quantified using FeatureCounts (v1.6.4) and used for downstream analysis.

### Differential gene expression analysis

Differential gene expression analysis was performed using the DESeq2 (v1.28.1) package in R. Significant differentially expressed genes (DEG) were filtered based on a p-adjusted (padj) < 0.05, and a log2 fold change (log2FC) greater than 0.50 or less than -0.50.

### Deletion of Lamin B1 in mouse splenic germinal centre B cells

All mice were on a C57BL/6 background using the *Cre*/*loxP* system, which engages in the *Cre*-mediated deletion of *LMNB1*. GC-targeted *LMNB1* deletion was accomplished by crossing the C57BL/6N-Lmnb1^tm1c(EUCOMM)Wtsi^/WtsiH (referred to as *Cγ1cre^+/-^;Lamin B1^fl/fl^*) strain, obtained from EMMA (EM:10615), and B6.129P2(Cg)-*Ighg1^tm1(cre)Cgn^*/J (referred to as *Cγ1cre^+/^*^-^), kindly provided by Dr. Dinis Calado (Crick Institute, UK) (Casola, Cattoretti et al., 2006). *Cre* recombinase activation was induced by the stimulation of the immune response after an intraperitoneal (IP) injection of 100 μL ImjectTM Alum Adjuvant with 100 µg of NP-CGG (Chicken Gamma Globulin) Ratio 10-19 (N-5055B-1, 2 B Scientific) at a concentration of 0.5 mg/mL. *Cγ1-Cre* mice were used as heterozygous for the *Ighg1*<tm1(cre)Cgn> gene, and the LaminB1^fl/fl^ mouse strain was homozygous for Lmnb1^tm1a(EUCOMM)^/WtsiH. All control and experimental mice were randomised among littermates between 8-12 weeks old. All experiments involving mice were performed under the Queen Mary of London Veterinary Oversight with the UK Home Office authorisation.

### Flow cytometry of B cell membrane surface markers to assess B cell differentiation

To label cell surface B cell differentiation markers, a single-cell splenocyte suspension was prepared by disaggregating fresh spleen tissue using a 70 μm strainer and resuspending the cells in PBS (10% FBS). Red blood cells were lysed with 1x RBLB for 5 min at RT and washed with ice-cold PBS (2% FBS). Live cells were washed, stained with fluorochrome-labelled antibodies (see Resource Table), and analysed using an LSR Fortessa analyser (BD Biosciences, Oxford, UK). Single strains and FMOs were used to set up the compensation parameters and gating strategies, respectively. At least 10,000 cells were analysed per sample.

### Immunohistochemistry

Paraffin-embedded samples were dewaxed twice in 100% xylene for 3 min, followed by incubation in 100% ethanol for 3 min, and twice in 100% ethanol (3% H_2_O_2_) for 3 min to block endogenous peroxidases. Prior to the antigen retrieval step, the samples were successively placed in 100% ethanol and deionised H_2_O for 3 min. Antigen retrieval was performed by boiling the samples at the maximum temperature in a pressure cooker in the presence of 1 × antigen unmasking solution for 10 min. Slides were then placed in 1x DAKO wash buffer and incubated with primary and secondary antibodies and the VIP HRP Detection Kit (see Resource Table). Haematoxylin was used as the nuclear stain. The dehydrated slides were mounted and scanned with a digital slide scanner (NanoZoomer S210, Hamamatsu C13239-01) at 20X original magnification. Multiplex immunohistochemistry of the same tissue section was performed by stripping the previous staining and performing the protocol described above with the following antibody. At least four randomly selected GCs per spleen from a minimum of five mice were selected and used for the analysis. Raw images were processed and analysed using QuPath (v0.2.0) and Fiji (v2.0.0).

### Immunofluorescence

For immunofluorescence (IF) imaging, cells were washed with ice-cold phosphate-buffered saline (PBS) and spun onto a poly-L-lysine-coated slide for 5 min in a Shandon Cytospin centrifuge (#22004). Immediately after centrifugation, the cells were fixed in 4% paraformaldehyde (PFA) for 15 min and permeabilised with 0.2% Triton-X/PBS. Slides were then blocked with 3% bovine serum albumin (BSA)/PBS-T (0.2% Tween-20) for one hour and incubated with the primary antibody at specifically optimised concentrations and time durations according to the Resource Table. Slides were then washed with PBS-T and incubated with secondary fluorescently conjugated antibodies and DAPI at a concentration of 1 µg/mL for 45 min in the dark. Slides were mounted using ProLong Gold, and images were taken with a Nikon Ci-L fluorescent microscope or a confocal Zeiss LSM550/LSM710 microscope. Raw images were analysed using ImageJ software for the quantification of γ-H2AX and RAD51 foci, and 53BP1 foci were counted using the MATLAB FoCo script. The colour balance was adjusted using the same settings across allexperimental conditions.

### In-suspension breaks labelling in situ and sequencing (sBLISS)

To generate the genome-wide DSB profile, the sBLISS methodology was adapted in BL2 cells (control and shLMNB1) and FACS-sorted mouse splenocytes (GC B cells and Naïve B cells) after shLMNB1-mediated depletion and immunisation, respectively. BL2 cells were used in triplicate and mouse splenocytes were obtained from a minimum of four mice (Cγ1Cre^+/-^ and Cγ1Cre^+/-^ ; Lmnb1^fl/fl^) from two independent immunisations with 100 μg NP-CGG for ten days. Prior to cell sorting, we enriched the GC B cell population using the Germinal Centre B Cell MicroBead Kit, according to the manufacturer’s protocol. The sBLISS template was generated as previously described (Bouwman et al., 2020). Libraries were analysed using DNA Tapestation 4200 and sequenced as single-end (1 × 75 bp) reads on the NextSeq platform.

### Pre-processing of sequencing data

Data analysis was performed using a publicly available script created specifically for the sBLISS analysis. Raw FASTQ files were merged, and reads were filtered according to the associated sample barcode (see Table 1), allowing for a maximum of one mismatch in the barcode sequence. Prefixes in each read were trimmed and sequences were aligned to the hg19 or mm10 reference genome using BWA-MEM (v0.7.17) and bedtools (v2.30.0). Aligned reads with a mapping quality score ≤ 60 and PCR duplicates marked by SAM tools (v1.9) were filtered out. The generated BED files containing unique DSB locations were used for downstream analysis.

### Hotspot detection

The detection of sBLISS hotspots Macs2 (v2.1) was used to call peaks from the BED files of UMI-DSBs (no background model, no cross-correlation between strands around the hotspots, shift reads by -100 bp, and extend by 200 bp). DSB hotspots were defined as the peaks identified by Macs2 with a q-value < 0.05, fold enrichment > 4, and pileup > 10. Circus plots were used to visualise the unique sBLISS hotspots per sample.

### sBLISS and RNA-seq data integration

Significant DEGs from the shLMNB1 knockdown analysis (adjusted p-value < 0.05, log₂FoldChange > 1) were annotated with genomic coordinates using the EnsDb.Hsapiens.v75 using R package. Both the DEG genes and the sBLISS DSB sites were converted to GRanges objects, harmonised to the hg19 reference genome, and analysed for overlap.

### sBLISS and DLBCL common mutated genes analysis

We assessed whether DLBCL driver loci coincide with Lamin B1–dependent DSB hotspots by intersecting MACS2 peaks unique to shLMNB1 with curated DLBCL gene intervals. Hotspots (narrowPeak) were imported with rtracklayer, converted to UCSC chromosome style, and restricted to autosome. Overlaps were computed with GenomicRanges::countOverlaps, defining a gene as hotspot-positive if ≥1 shLMNB1-unique peak intersected its interval. Genes mutated in DLBCL patients according to the publicly available curated gene list generated by Ryan Morin Lab.

### Discovery of *de novo* motifs and analysis

To identify DNA *de novo* motifs enriched around Lamin B1-associated DSB hotspots, we used MEME Suite (v5.5.8). Sequences ±100 bp flanking the summit of high-confidence DSB peaks were extracted and analysed with motif widths constrained between 6 and 12 bp, and the top 5 motifs were retained. The top five most significant motifs identified by MEME were used for further analysis. The R packages universalmotif, ChIPseeker, and GenomicRanges were employed to scan for all motif occurrences within the identified peak regions of the *Mus musculus* genome (mm10 assembly). Genomic features (3’UTR, 5’UTR, CDS, exons, intergenic, introns, promoter and TSS) were hierarchically annotated, and associated genes were identified. To identify biological processes potentially affected by motif-associated DNA damage, we extended the genomic coordinates of *de novo* discovered motifs by ±3 kb and mapped them to the nearest TSS. This yielded a list of genes located in proximity to the motif hotspots. ClusterProfiler was used to perform a Gene Ontology (GO) enrichment analysis on the associated genes to infer the biological pathways linked to the discovered motifs.

The *de novo* motifs were also matched to known transcription factors (TF) motifs using Tomtom (MEME Suite) with the following parameters: -verbosity 1 -min-overlap 5 -mi 1 -dist pearson - evalue -thresh 10.0 -time 300. We queried against two motif collections: the Uniprobe mouse database and the JASPAR 2024 CORE vertebrates non-redundant database. Matches were ranked by Pearson correlation and minimum information content overlap of at least five positions. To contextualise the biological relevance of matched TFs, we utilised the CistromeDB, querying publicly available ChIP-seq data from different cell types to evaluate TF binding site enrichment near our input regions. GIGGLE similarity scores were computed, and enrichment analysis against reference epigenomic datasets was performed.

### Clinicopathological analysis of *LMNB1* expression and GEP in CLL and DLBCL

For CLL gene expression analysis with respect to LMNB1 expression, we downloaded a dataset from patients with CLL (REACH trial GEO: GSE52811). Normalisation, cluster analysis, and subtype classification were conducted as previously described (Bloehdorn et al., 2021). *LMNB1* expression was assessed in the GI subtype (highest frequency of *TP53* alterations, shortest progression-free survival (PFS) in *TP53* WT cases), in comparison to the (I)EMT-L subtype (lowest frequency of *TP53* alterations, most prolonged PFS). For DLBCL, we utilised a publicly available dataset from TCGA (phs001444) to assess *LMNB1* expression and the clinical outcomes of 481 DLBCL patients (Schmitz et al., 2018). Raw count files and clinical data were obtained from TCGA and the previously published DLBCL study (Schmitz et al., 2018), respectively. Log2-transformed counts per million (Log2CPM) were used for GEP and clinical evaluation of *LMNB1* expression. Kaplan-Meier survival analysis for PFS was conducted based on *LMNB1* expression levels in DLBCL patients. The group cutoff was set based on the quartile or median *LMNB1* expression. Ethical approval was obtained from the East London & The City Health Authority Local Research Ethics Committee (Reference number 10/H0704/65), and written informed consent was obtained in accordance with the Declaration of Helsinki.

### Study approval

All animal studies were conducted in compliance with the UK Home Office-approved licences (Animals (Scientific Procedures) Act 1986 and the EU Directive 2010). Animals were maintained in local facilities, and the local ethical committees under the Home Office license P68650650 approved experiments.

### Statistical analysis

Generated data were obtained from independent biological replicates, as indicated in the figure legend or the corresponding methods section. For IHC, IF, and western blotting analysis, raw images were processed using Fiji and QuPath. Foci counting was performed using the Fiji or FociCounter algorithm in MATLAB (MathWorks). Survival analysis was performed in R using the ggsurvplot package. Numerical data were analysed and plotted in Prism v8.3 (GraphPad Software, Inc.) or R. P values were calculated using a two-tailed unpaired t-test unless stated otherwise.

## Declarations

### Data and materials availability

All data needed to evaluate the conclusions in the paper are presented in the paper and/or the Supplementary Materials. Sequencing files were deposited in the Sequence Read Archive Project PRJNA892748.

### Author contributions

Conceptualisation, FF, MCS, AB, and TK; Methodology, FF, MCS, AB; Investigation, FF, MCS, JB, JW, Software, MCS, JW, JB, FF; Formal Analysis, FF, MCS, JW, JB; Writing – Original Draft, FF, MCS; Writing – Review and Editing, FF, MCS, TK, JG, KC, MH; Funding Acquisition, AB; Resources, JG, CC, MH, AB; Supervision, TK, MCS and AB.

## Acknowledgments

We thank Findley Copley, Andrew Clear, Nicolas Peschke, and Marton Gelleri for their technical assistance.

---

**Supplementary Figure 1.**
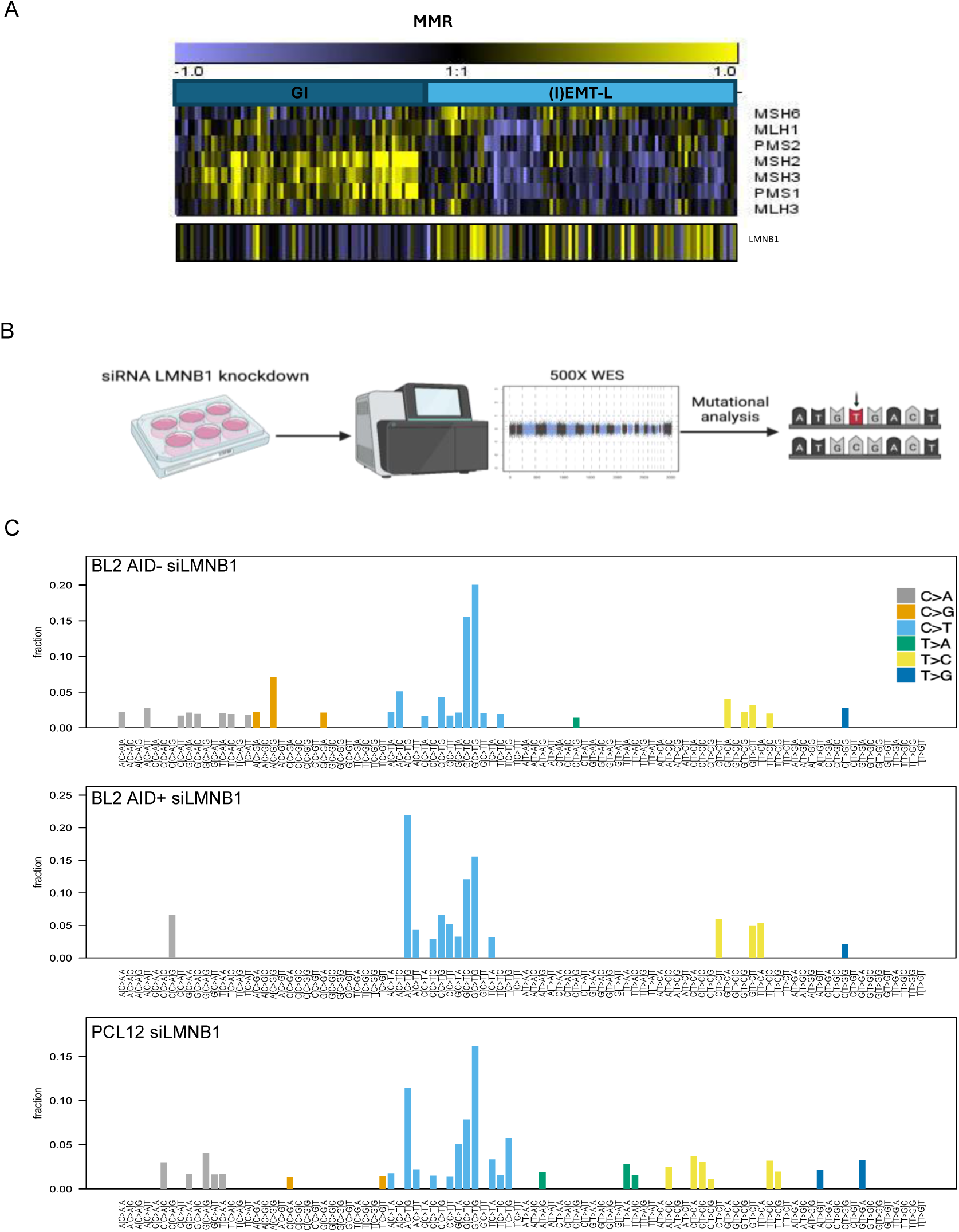
Decreased Lamin B1 is associated with CLL genomic instability and mutagenesis in malignant B cells. Decreased Lamin B1 is associated with CLL genomic instability and mutagenesis in malignant B cells. **(A)** Heatmap showing GEP of DNA mismatch repair (MMR)-associated genes in GI and (I)EMT-L CLL subtypes (*n* = 173). **(B)** Schematic representation of the WES experiment to deconvolute the mutational pattern in siLMNB1 lymphoma cells. **(C)** Representative analysis of single base substitutions (SBS) detected in Lamin B1 knockdown cell lines (BL2 AID-, BL2 AID+, PCL12), confirming the enrichment in C>T substitution.

**Supplementary Figure 2.**
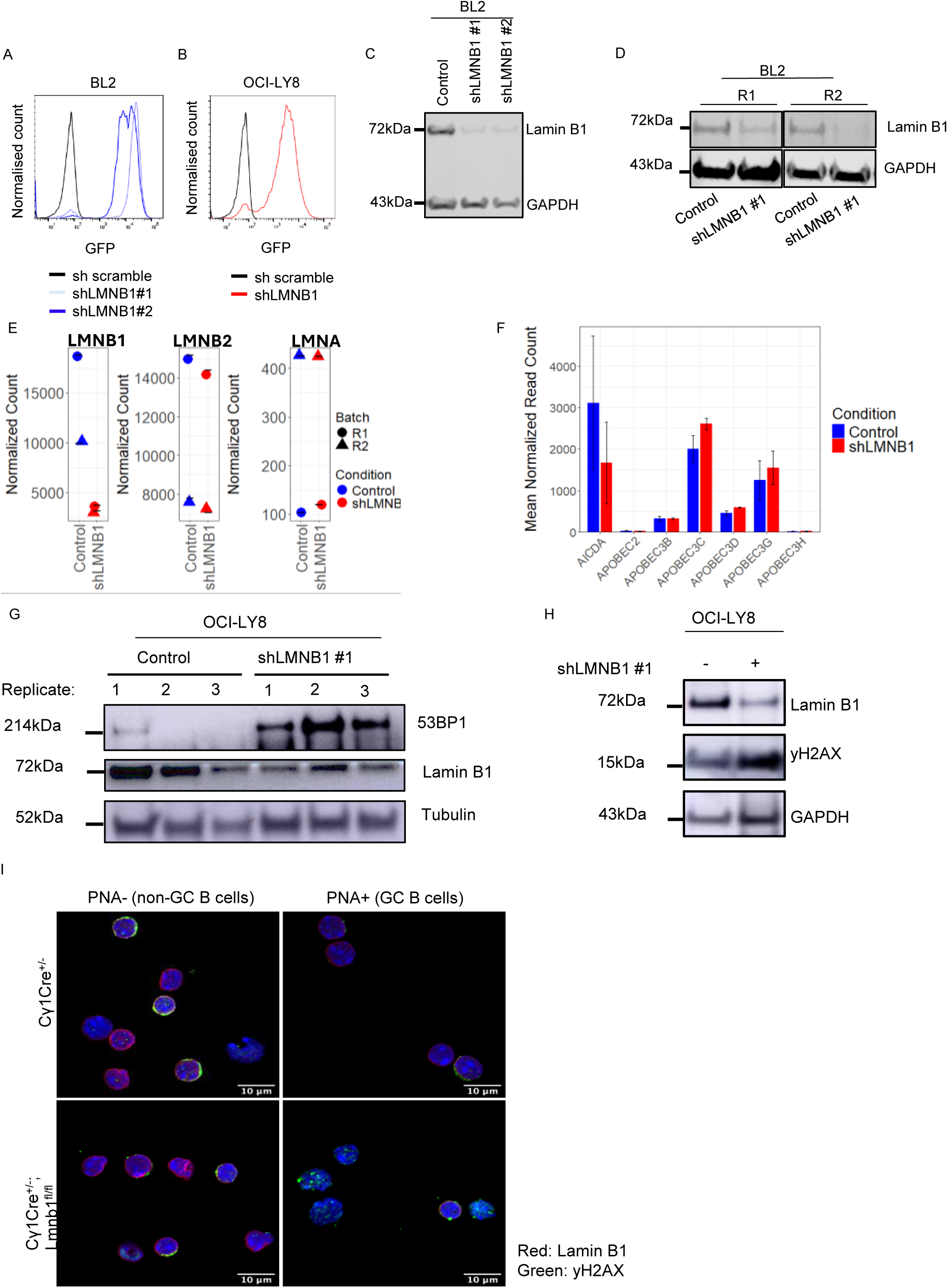
*In vitro* Lamin B1 reduction translates into increased GI, unrelated to increased expression of AICDA or APOBEC genes. **(A)** Flow cytometry analysis of GFP expression following shLMNB1 induction in BL2 cells. **(B)** Flow cytometry analysis of GFP expression following shLMNB1 induction in OCI-LY8 cells. **(C)** Western blot showing Lamin B1 expression in control and shLMNB1-expressing BL2 cells. **(D)** Western blot confirming reduced Lamin B1 expression in RNA-seq replicates R1 and R2. **(E)** Normalised RNA-seq expression levels of LMNA, LMNB1, and LMNB2 comparing control and shLMNB1 conditions. **(F)** Histogram depicting expression levels of AICDA and APOBEC family members in control versus shLMNB1 conditions, derived from RNA-seq data. **(G)** Western blot showing the expression of 53BP1 and Lamin B1 in OCI-LY8 cells across three replicates. **(H)** Western blot analysis of Lamin B1 and phospho-γH2AX levels in OCI-LY8 cells following shLMNB1 induction **(I)** Representative immunofluorescence images of PNA⁻ and PNA⁺ cells isolated from Cγ1Cre^⁺/⁻^ and Cγ1Cre^⁺/⁻^; Lmnb1^fl/fl^ mice.

**Supplementary Figure 3.**
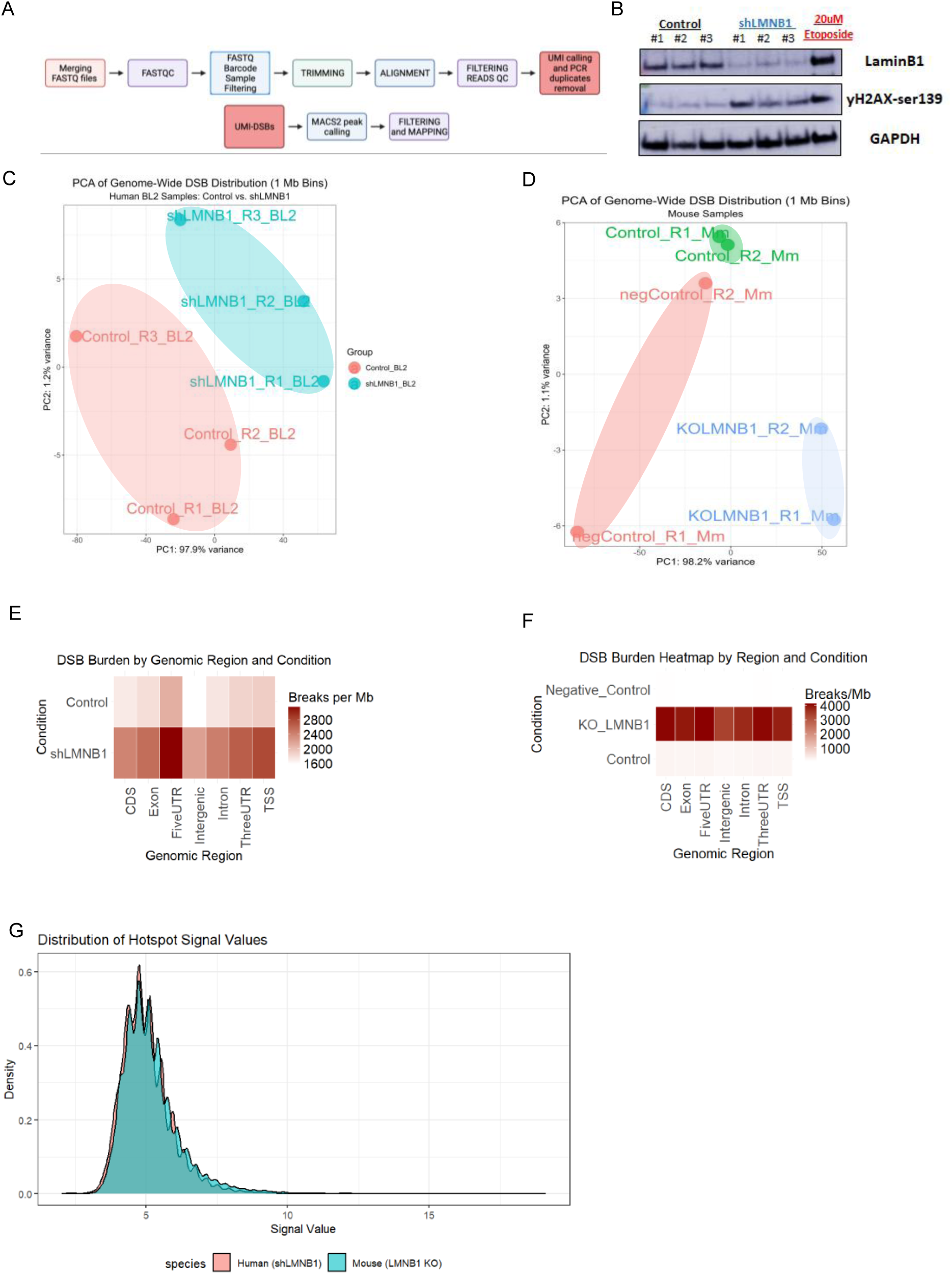
DNA break profiling reveals that *LMNB1* reduction induces genome-wide genomic instability. (**A**) Schematic overview of the bioinformatic pipeline for sBLISS data analysis. **(B)** Western blot showing Lamin B1 and phospho-γH2AX expression levels before the sBLISS experiment. **(C-D)** PCA plot of genome-wide DSB distribution (1Mb Bins) in (**C**) human BL2 samples and (**D**) mouse samples. Heatmap displaying the DSB burden per genomic region in (**E**) BL2 cells and (**F**) *in vivo* GC B cells and non-GC B cells across experimental conditions. (**G**) Histogram showing the distribution of hotspot signal values in human shLMNB1 and mouse LMNB1 knock-down GC B cells.

**Supplementary figure 4.**
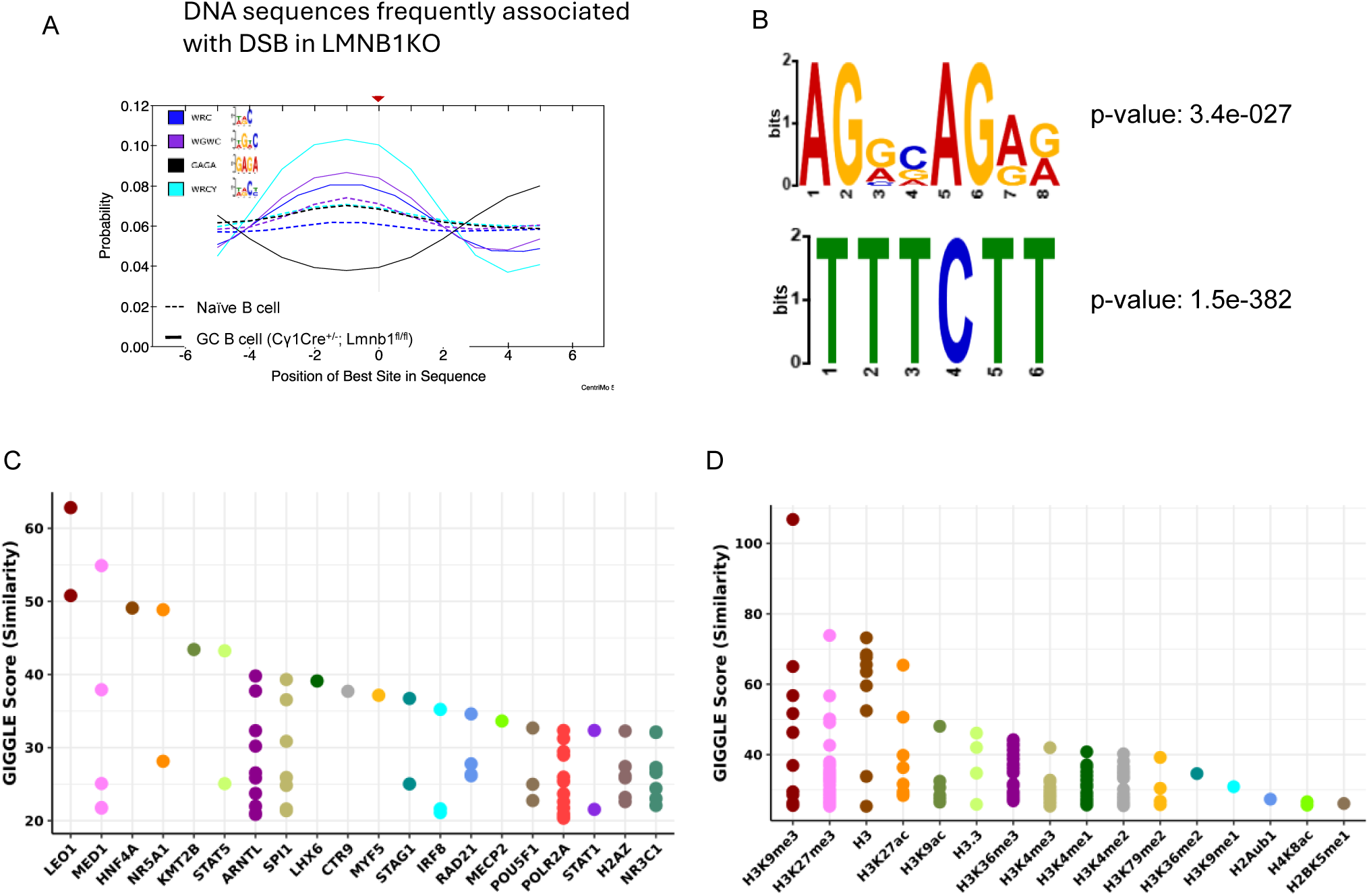
Lamin B1 associates DSB hotspots with chromatin regulators. **(A)** AID hotspots (WRC, WRCY, and WGCW) location probability associated with DSBs in Naïve B cells compared to *Cγ1cre^+/-^;Lmnb1^fl/fl^*GC B cells. (**B**) Top 4 and 5 enriched *de novo* motifs in *Cγ1cre^+/-^; Lmnb1^fl/fl^* GC B cells. (**C-D**) Cistrome analysis with GIGGLE scoring quantified the association between unique DSB hotspots, transcription factors (**C**) and histone modifications (**D**) in *Cγ1Cre⁺/⁻;Lmnb1^fl/fl^* mice.

**Supplementary figure 5.**
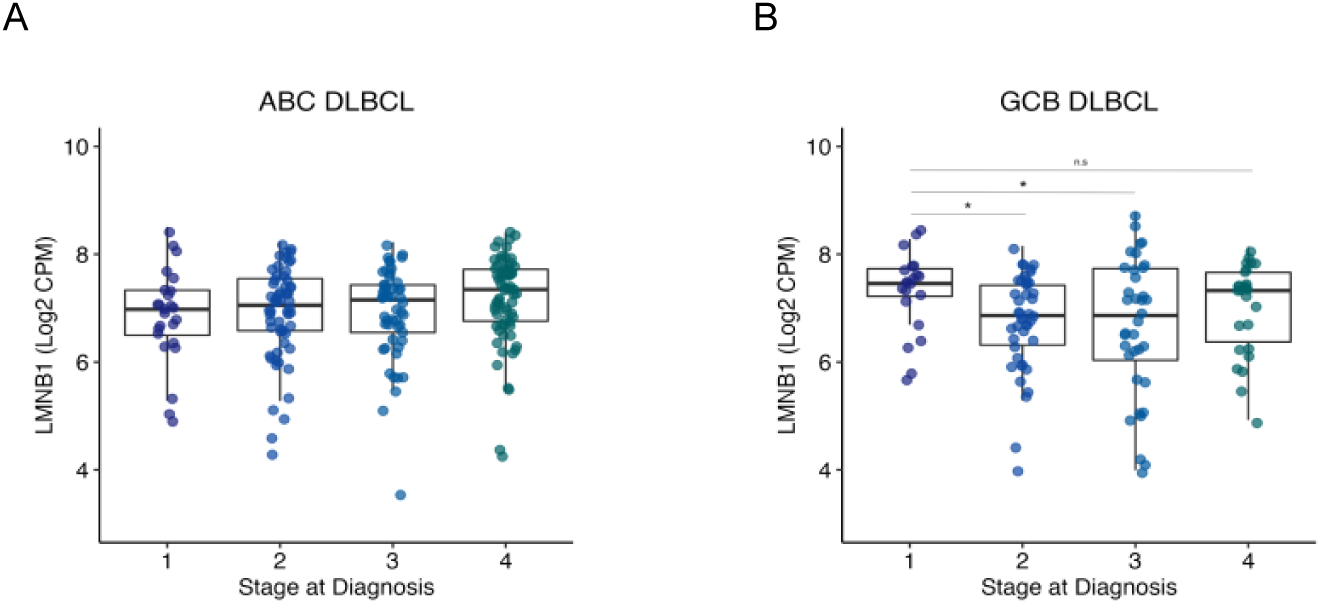

**Supplementary figure 6.** Decreased *LMNB1* expression is associated with worse clinical outcomes in DLBCL patients. (**A-B**) Boxplot showing *LMNB1* expression across diagnostic stages in (**B**) ABC DLBCL subtypes (N=205) and (**C**) GCB DLBCL subtypes (N=126).

## Supplementary Resource Table

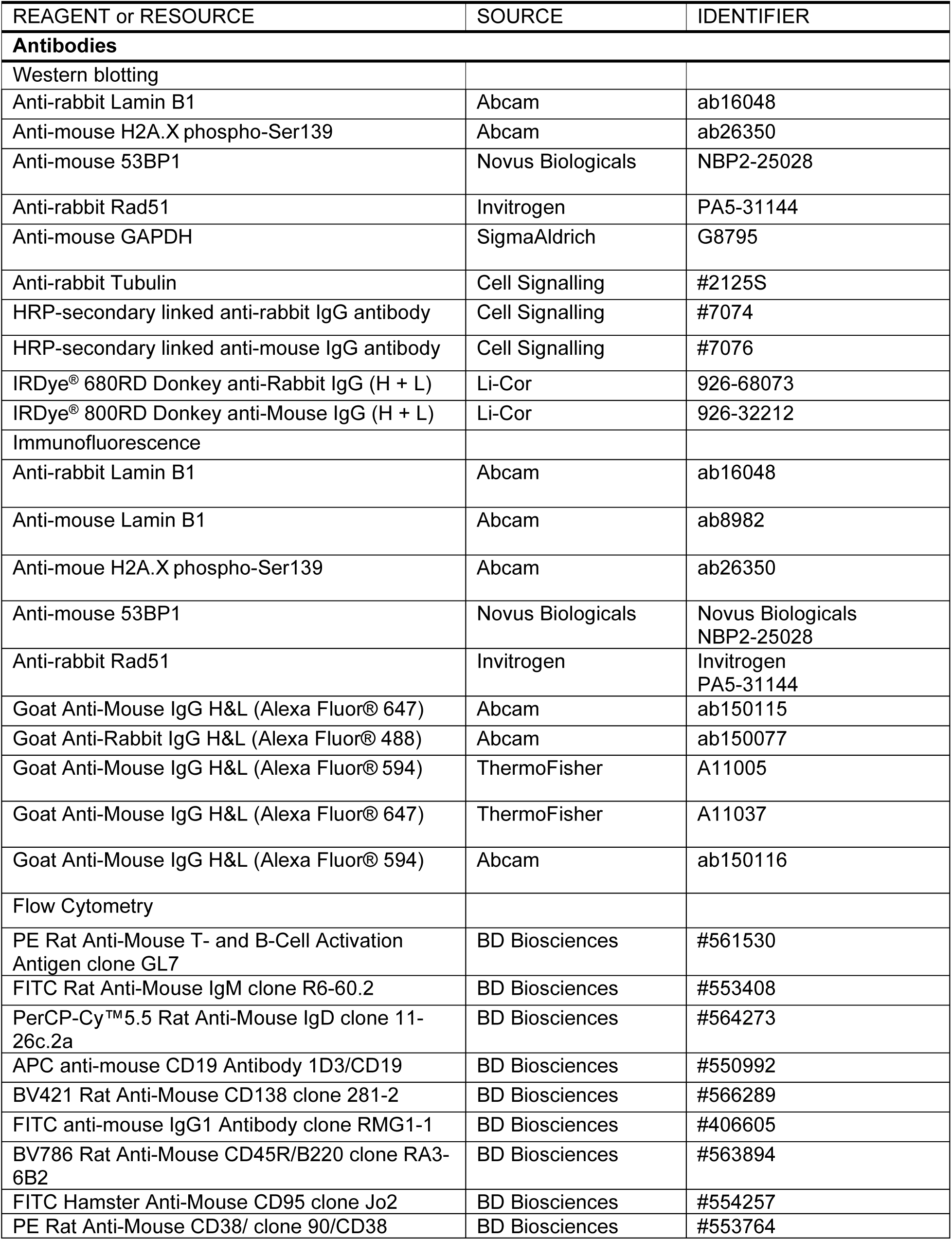
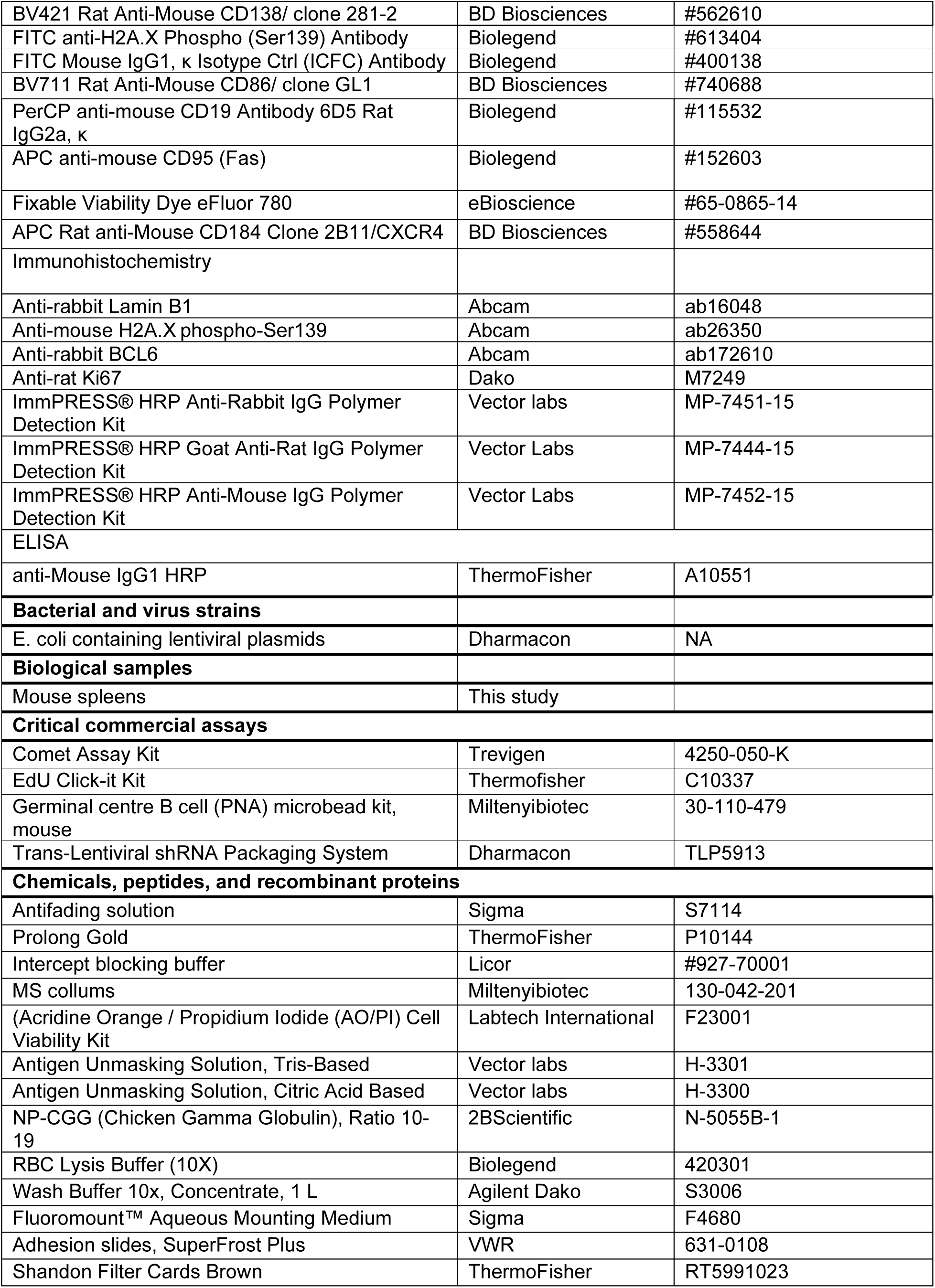
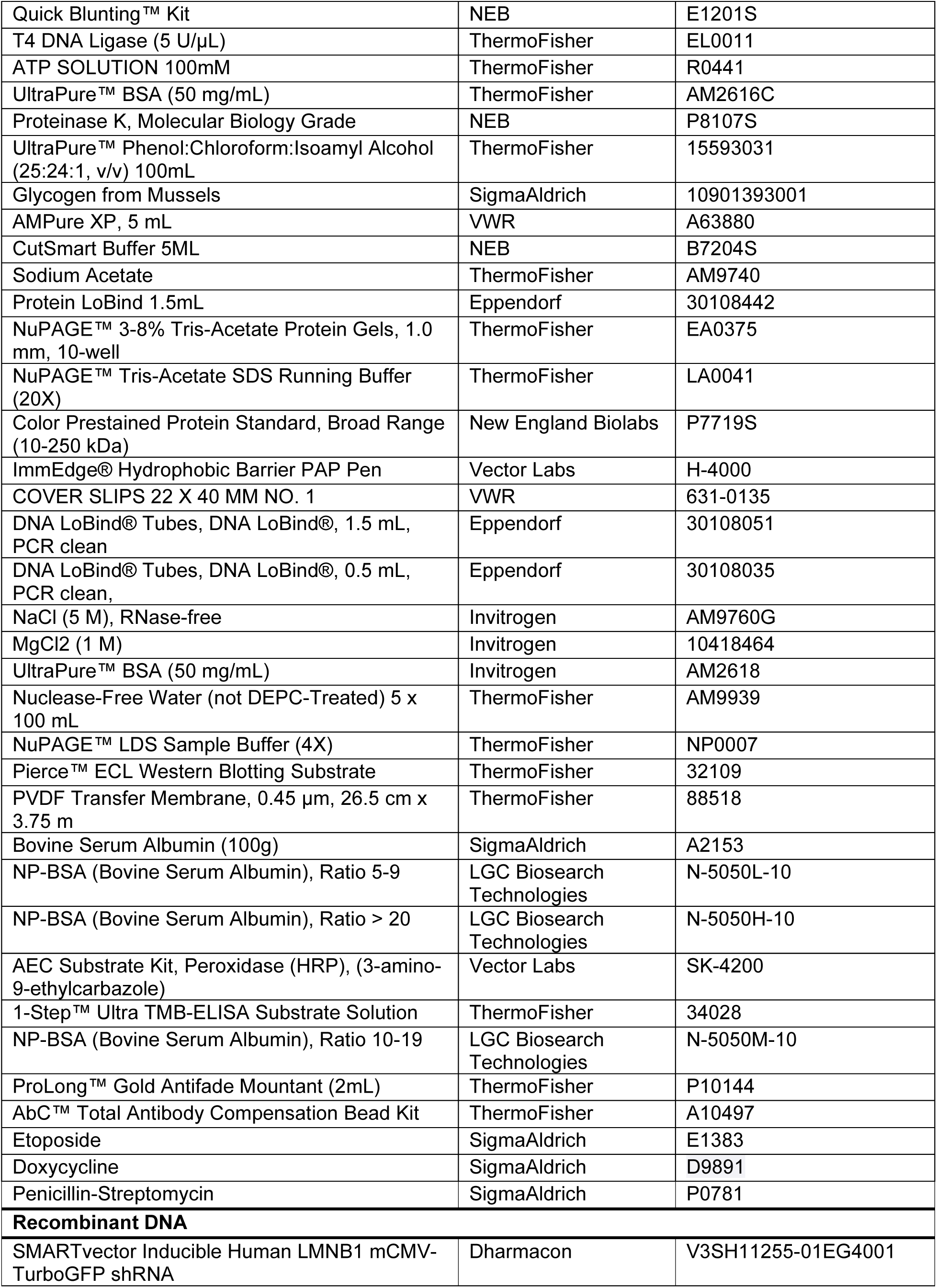
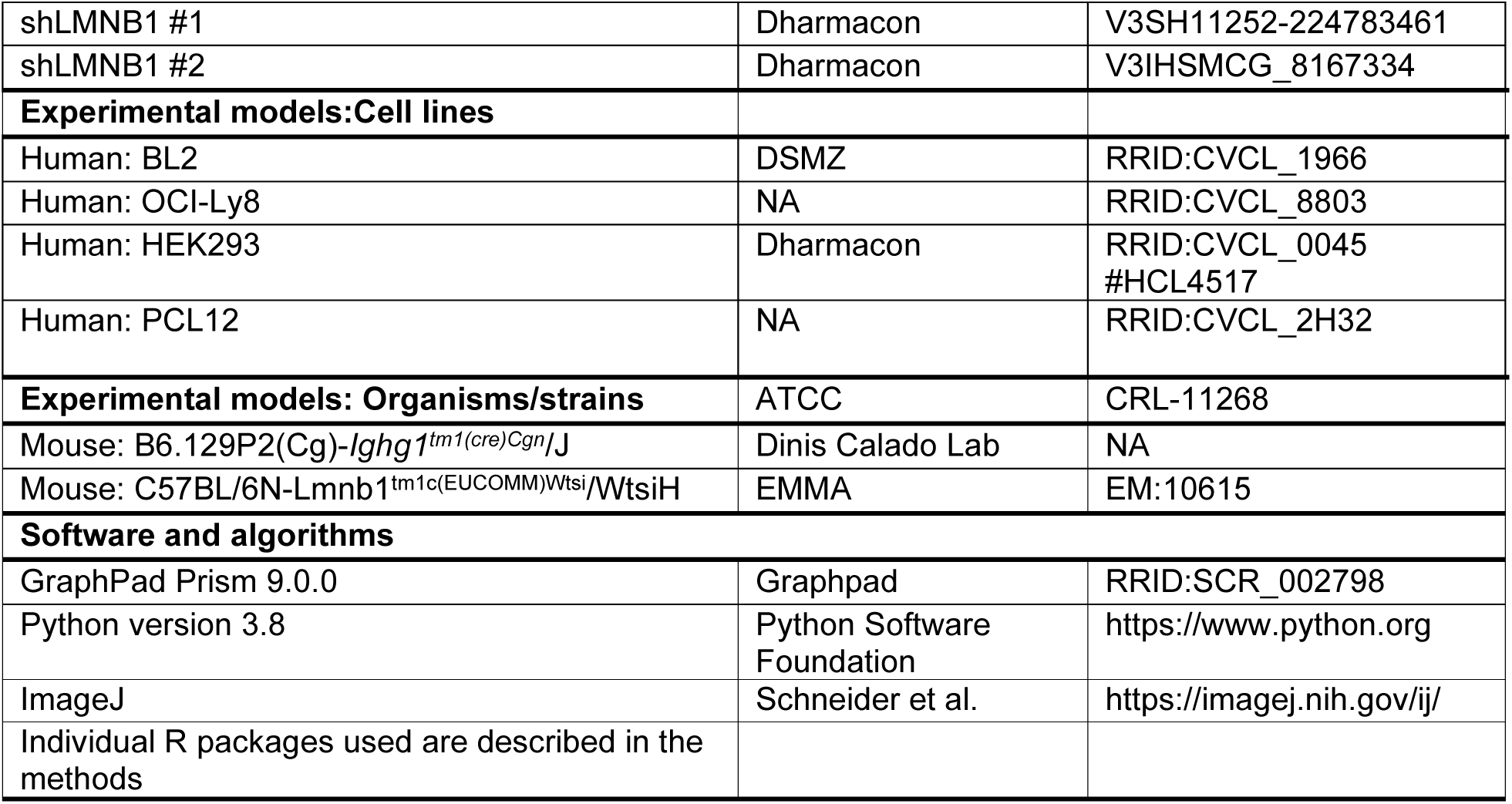

